# Unraveling Resistance and Thresholds in Phage Therapy Using *Staphylococcus aureus* In Vitro Models

**DOI:** 10.1101/2025.04.25.650594

**Authors:** Kévin Royet, Leslie Blazere, Emilie Helluin, Ludivine Coignet, Lucile Plumet, Mélanie Bonhomme, Camille Kolenda, Romain Garreau, Florent Valour, Mathieu Medina, Frédéric Laurent, Sylvain Goutelle

## Abstract

**Background:** Phage therapy is a promising approach against multidrug-resistant *Staphylococcus aureus*, but its clinical development is limited by gaps in pharmacokinetic/pharmacodynamic (PK/PD) understanding, especially concerning phage self-replication and resistance dynamics.

**Methods:** We combined *in vitro* time-kill assays with PK/PD modeling to study three anti-staphylococcal phages (V1SA019, V1SA020, V1SA022) against two *S. aureus* strains (SH1000 and USA300). A system of differential equations captured the co-dynamics of susceptible, resistant and infected bacteria and free phages. Nonlinear mixed-effect modeling quantified parameter variability. Resistance emergence was monitored through phenotyping and whole-genome sequencing of resistant clones.

**Results:** Phage–bacteria interactions followed a predator–prey pattern, with early bacterial growth, rapid phage amplification, and subsequent bacterial collapse. However, resistant subpopulations emerged, regrew over time, in a multiplicity of infection (MOI) dependent way. The model accurately described bacterial and phage dynamics and estimated kinetic parameters including adsorption and burst size. Proliferation and inundation thresholds varied by strain and phage. All resistant clones harbored mutations in genes involved in teichoic acid biosynthesis, with associated growth defects. Simulations demonstrated that only phage doses exceeding both susceptible and resistant bacterial inundation thresholds fully suppressed regrowth.

**Conclusion:** This study provides a quantitative framework for understanding phage–*S. aureus* co-dynamics and resistance emergence. It emphasizes the importance of considering both proliferation and inundation thresholds when designing phage dosing regimens. These findings inform the rational development of phage therapy and support translation toward *in vivo* and clinical applications.

## Introduction

The emergence of antibiotic resistance is one of the major public threats requiring the development of new antibiotics or alternative therapeutic strategies. Therapy using bacteriophages (or phages) represents a source of considerable hope [1], [2]. Phage therapy has been developed and used for decades in Eastern countries, while in Western countries it is considered that clinical evidence is currently insufficient to be approved by health authorities. In those countries, phage therapy is only used on a compassionate basis when conventional treatments are ineffective. However, phage therapy is promising and was reported to be life-saving in several cases [3].

The limited availability of pharmacokinetic and pharmacodynamic (PK/PD) data, combined with the underdevelopment of mathematical models capable of integrating these parameters, hinders the generation of predictive efficacy data of phage therapy. The PK/PD of phages is very different from that of antibiotics and more complex, since phages can self-replicate in the presence of their target bacteria [4]. The co-dynamics of phage and bacterial populations are characterized by specific thresholds effects. The proliferation threshold refers to the bacterial cells density above which the probability of a progeny phage replicating is greater than the probability of that phage being lost [4]. The inundation threshold corresponds to the phage density above which the probability of a bacterial cell being lysed by a phage is greater than the probability of that same bacterial cell dividing [5]. Quantifying those thresholds is crucial in phage therapy in order to define effective doses: the first ensures immediate action of the phages even without replication, while the second guarantees that local amplification of the phages is possible depending on the bacterial density. This allows the therapeutic strategy to be optimized according to the type and dynamics of the infection.

*Staphylococcus aureus* is responsible for various infections, including bacteremia, sepsis, pneumonia, endocarditis and bone and joint infections (BJI). Resistance of *S. aureus* to antibiotics is widespread, with methicillin resistant strains being clinically significant [6]. Because of its ability to form biofilms, this pathogen is also difficult to eradicate in device-associated infections. For those reasons, *S. aureus* is classified as high priority for research and development of new therapeutics by the World Health Organization and is a target of interest for phage therapy [7].

The objectives of this study were (i) to describe and quantify the *in vitro* co-dynamics of three candidate anti-staphylococcal phages (V1SA019, V1SA020 and V1SA022) and two reference methicillin susceptible (SH1000) / resistant (USA300) *S. aureus* strains; and (ii) to build a robust mathematical model describing these co-dynamics and that can generate predictive data of phage efficacy.

## Materials and methods

### Experimental method

#### Bacterial culture and phage propagation

We used two reference strains of *S. aureus*, SH1000 [8] and USA300 LAC strain [9], as well as three phage candidates, V1SA019, V1SA020 or V1SA022 [10].*S. aureus* strains were grown weekly on Columbia Agar supplemented with 5% Sheep Blood (BioMerieux, Marcy-l’Etoile, France) for 20 h at 37 °C. For culture, one colony was picked with an inoculation loop and suspended in cation adjusted Mueller Hinton (caMH) (Becton Dickinson, Franklin Lakes, NJ, USA), then incubated at 37 °C for 20 h at 180rpm. *S. aureus* bacteriophage V1SA019, V1SA020 or V1SA022 were routinely propagated as previously described [10]. Briefly, susceptible bacterial strain in exponential phase and phages were mixed at MOI of 10^−3^ phage per bacteria in 10 mL of Tryptic Soy Broth (TSB, BD, Franklin Lakes, NJ, USA). After 24 h of incubation, the obtained phage lysates were filtered using a 0.22 µm syringe filter and phage titers were measured by spot test assays. For spot test assay, 600uL of overnight culture of a susceptible strain was mixed with 3.5% caMH agar melted and kept at 55°C and then poured in an empty 120-mm-diameter petri dish. Finally, phages were serially diluted in PBS, and 5 μL of each dilution were spotted onto plates, which were then incubated for 24 hours at 37 °C prior to plaque-forming unit (PFU) enumeration.

#### Bacteriophage decay

To determine the free phage decay rate in the absence of bacteria, V1SA019, V1SA020 or V1SA022 were added to 25 mL of pre-warmed caMH in 50 mL conical tubes at a final concentration of 10^1^ PFU mL^-1^, 10^3^ PFU mL^-1^ or 10^5^ PFU mL^-1^ and incubated at 37 °C with shaking at 180 rpm. Samples were collected at 3 h, 6 h, 24 h and 48 h and bacteriophages were enumerated using double layer plaque assay [11]. Briefly, 100μL of a phage lysate was mixed with 300μL of bacterial culture in stationary phase and held for 10min at room temperature. Then, 10 ml of 3.5% caMH agar melted and kept at 55°C was added, and, after mixing, the contents of the tube were immediately poured onto a 90-mm-diameter petri dish of caMH agar (BioRad, Hercules, CA, USA). Three independent experiments were performed.

#### *In vitro* time-kill experiments

Four series of time-kill experiments were conducted. The effect of the three phages (V1SA019, V1SA020 or V1SA022) was evaluated on the SH1000 strain. In addition, V1SA019 was also tested against the USA300 LAC strain. This allowed to study the variability of the phage-bacteria dynamics between phages and bacterial strains. Optical density at 600nm (OD_600nm_) of overnight *S. aureus* liquid cultures was measured and an aliquot was added to 25 mL pre-warmed caMH in conical tube (Falcon, Le Pont de Claix, France, 50 mL) to a final concentration of 10^3^ colony-forming units (CFU) mL^-1^ and incubated at 37 °C under microaerobic condition with shaking at 180 rpm. After 1 h of lag phase, bacteriophages V1SA019, V1SA020 or V1SA022 were added to a final concentration of 10^1^ plaque-forming units (PFU) mL^-1^, 10^3^ PFU mL^-1^ or 10^5^ PFU mL^-1^ to give an approximate multiplicity of infection (MOI) of 0.01, 1 or 100. Samples were taken every 2 h for 24 h, serially diluted and plated on caMH agar plates for bacterial enumeration. Undiluted samples were also spread on caMH agar to lower down the detection limit to 4 CFU mL^-1^. Each aliquot was then centrifuged for 30s at 20,000g to remove *S. aureus* cells and supernatant was diluted for bacteriophages enumeration by spot test or double layer plaque assays when applicable (i.e., when estimated PFU mL^-1^ < 2.2 x 10^2^). At the end of the experiment, 1mL of culture was stored at-80°C in culture medium with 20 % glycerol to allow subsequent isolation of bacteria. Experiments were performed using three different phage production batches infecting tree independent bacterial cultures for each MOI. All experiments were conducted in triplicates on different weeks.

#### PK/PD modeling

We used a population approach and nonlinear mixed effect modeling to describe the time course of bacteria and phage counts in all four sets of experiments. Time-kill, bacterial growth and phage decay data were analyzed. However, adsorption and burst-size data could not be included in the final model, as preliminary runs showed model instability and unrealistic estimates. Nevertheless, adsorption and burst size assays are described in supplementary methods and results are shown in Table S1, S2 and Figure S1, S2.

A model written in ordinary differential equations (ODE) was implemented in the Monolix software (version 2024, Lixoft, Antony, France) to describe the interactions between four populations: susceptible bacteria (S), resistant bacteria (R), bacteria infected by phages (I) and free phage viruses (V). The ordinary differential equations corresponding to this model were as follows (Equation 1):

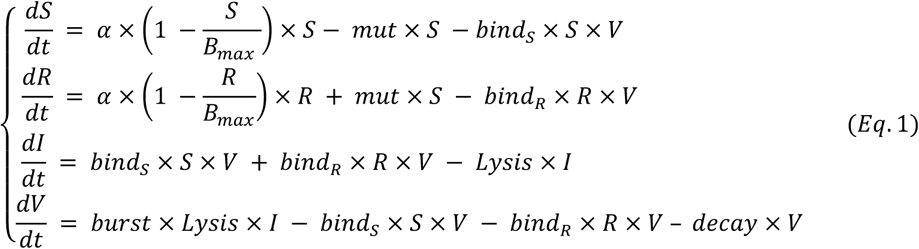

The eight parameters were:

α: bacterial growth constant; Bmax: maximal concentration of bacteria; mut: mutation rate of S to R bacteria; bind_S_ and bind_R_: binding rate constant of phages to S and R bacteria respectively; lysis: lysis rate constant of I bacteria; burst: number of free phages released upon I bacteria burst; decay: elimination rate constant of free phages

This model was inspired by previously published *in vitro* PK/PD studies [5], [12] and is schematically described in Figure 1. Inclusion of resistant bacteria was necessary to describe bacterial regrowth observed *in vitro*. For each parameter, both fixed (median, typical value) and random (inter-experiment variability) effect parameters were estimated, within a general framework as follows:

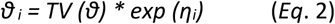

**Figure 1.**
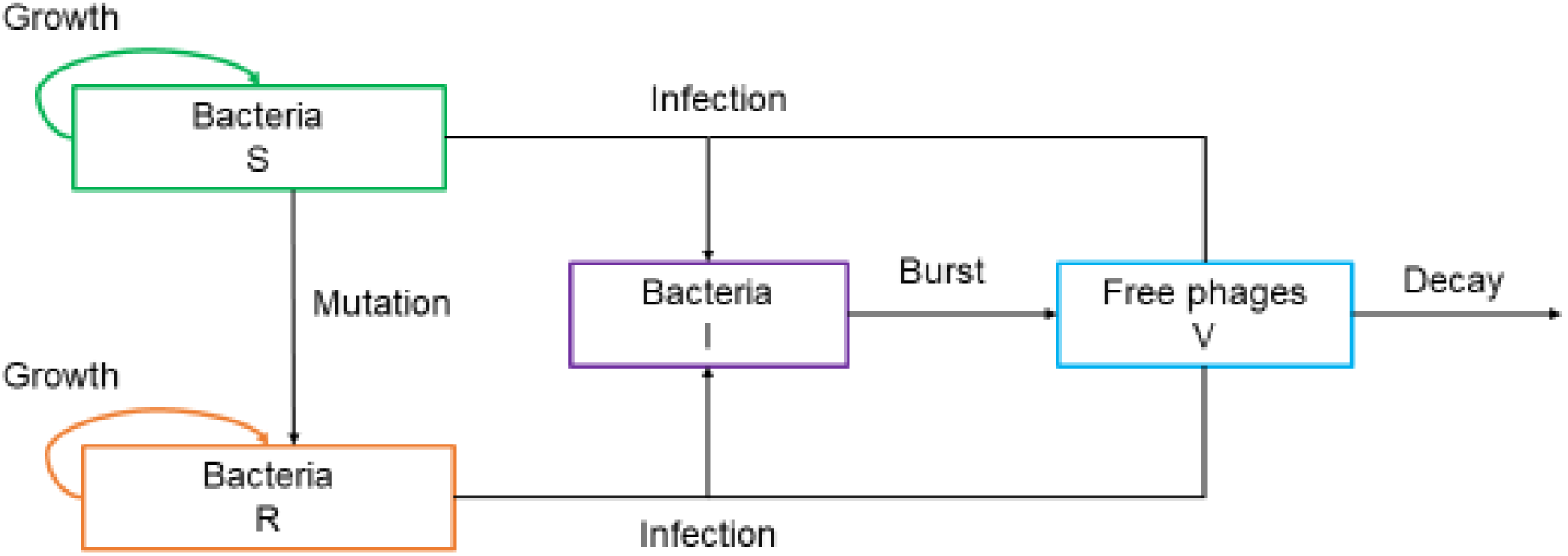
Structural model of the bacteria – phage co-dynamics

With

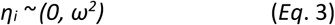

Where θi is the parameter value (e.g., bacteria growth rate, phage decay…) for an individual experiment, TV(θ) is the typical parameter value in the whole set of experiments, i.e. the median for a lognormal distribution (considered as a fixed effect in the nonlinear mixed effects model), and η_i_ is a random effect variable quantifying the variability around the typical value. Random effects were assumed to follow a normal distribution with mean of zero and variance ω². As a result of Equation 2, parameters (*θ_i_*) are assumed to follow a log-normal distribution.

Model outputs were coded as log_10_(S + R + I) and log_10_(V) for bacteria and virus, respectively, to facilitate comparison with observations. Several residual error models were evaluated, including additive, proportional and combined error models. The production batch was evaluated as a potential descriptor of inter-experiment variability.

Model evaluation was based on goodness-of-fit criteria including objective function (-2 x log-likelihood) and Akaike Information Criterion (AIC). Plots of observed versus model individual estimates were examined, as well as distributions of individual weighted residuals. Visual predictive checks based on 1000 simulations were performed as internal validation. Because proliferation and inundation thresholds are not primary parameters but combination of ODE parameters [5], they were not estimated at the population level but computed for each experiment after Bayesian estimation of individual parameters and summarized with descriptive statistics.

The proliferation thresholds of phages for susceptible and resistant bacteria (Prolif_S_ and Prolif_R_) were calculated as follows [5]:

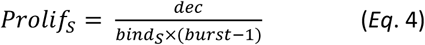

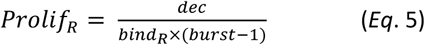

The inundations thresholds for susceptible and resistant bacteria (Inund_S_ and Inund_R_) were calculated as follows [5]:

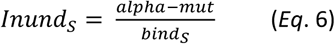

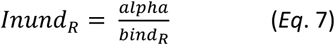

Finally, simulations were performed based on the final model for V1SA019 / SH1000 to study the influence of phage MOI on antibacterial effect.

### Liquid phage susceptibility test

From *S. aureus* clones isolated and grown on COS agar plates, a colony was resuspended in a 96-well plate containing 200 µL of caMH alone or with 10^6^ PFU mL^-1^ of phage. Plates were incubated at 37 °C using LogPhase 600 under agitation (Agilent BioTek, Santa Clara, USA), and OD_600_ measurements were taken every hour for 24 h. A clone was defined as resistant when its optical density at 600 nm after 24 hours of incubation exceeded 0.1, in contrast to the negative control (wild-type SH1000 inoculated in the presence of the phage), which showed no detectable growth under the same conditions.

### DNA extraction and sequencing

Bacterial DNA was extracted using the Maxwell® RSC Blood DNA kit (Promega, Madison, WI, USA), after an initial incubation with 40 μg lysostaphin (Sigma-Aldrich, Saint Louis, MO, USA Aldrich), 200 μg lysozyme (Sigma-Aldrich) and 200 μg RNaseA (Qiagen, Hilden, Germany) for 1 h at 37 °C, followed by incubation with 12 mAU (AU= Anson unit) of proteinase K (Promega) for 20 min at 60 °C. Whole genome sequencing was performed with NextSeq instrument (Illumina, San Diego, CA, USA) using a 150 bp paired-end protocol. Reference genome assembly was performed using SPAdes v3.15.5 [13] and Bakta v1.7.0 [14] was used for genome annotation. SNPs analysis was conducted with snippy v4.6.0 (https://github.com/tseemann/snippy). Indels were found with mummer4 v4.0.0 [15].

### Assessment of the fitness of phage-resistant clones

The relative fitness of phage-resistant clones was assessed by comparing their growth dynamics to that of the wild-type strain. Strains were pre-cultured overnight in caMH medium at 37°C, then diluted to an initial OD₆₀₀ of 0.01 in fresh caMH medium. Cultures were transferred to 96-well microplates and incubated at 37°C with continuous orbital shaking. OD₆₀₀ measurements were recorded hourly using a Tecan Infinite M200 Pro plate reader. Each condition was tested in four independent biological replicates, each comprising three technical replicates.

### Statistical analysis of experimental data

Statistical analyses and plotting were performed using GraphPad Prism, version 8.0.0 (GraphPad Software, San Diego, CA, USA). Variables were compared using the nonparametric Mann-Whitney U test. A p-value < 0.05 was considered significant.

## Results

### Phage decay

Without bacteria, phages declined linearly on the log-scale (Figure 2). The loss of phage viability over 24 h was low (mean ratio < 10 PFU.mL^-1^ between 0 - 24 h) for all the concentrations tested, although significant for inoculum values of 10^1^ PFU mL^-1^ and 10^5^ PFU mL^-1^ of V1SA019, V1SA020 and V1SA022. Over 48 h, phage decay was larger, with mean ratios of phage decay between 0 h and 48 h of 5.4, 3.6, 3.2 for V1SA019; 7, 27, 39 for V1SA020; 9, 34, 43 for V1SA022 for inoculum of 10^1^ PFU.mL^-1^, 10^3^ PFU.mL^-1^ and 10^5^ PFU.mL^-1^, respectively.

**Figure 2.**
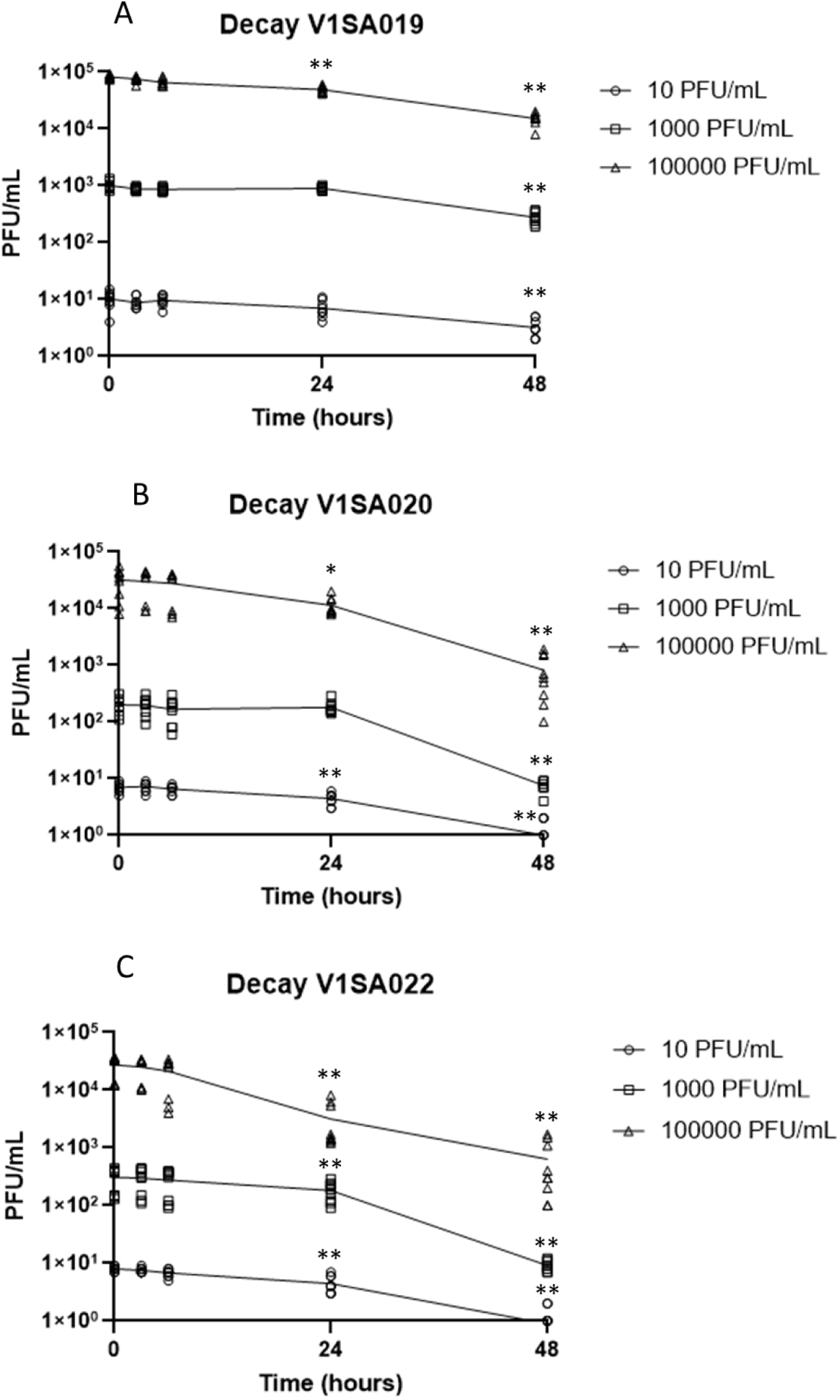
Phage decay of V1SA019 (A), V1SA020 (B), V1SA022 (C). Each symbol represents an experimental point. Connecting line represents the median. Statistical analysis compared two time points via Wilcoxon matched pairs signed rank test. *: p-value < 0.05; **: p-value < 0.005.

### *In vitro* time-kill studies

Phage-bacteria kinetic experiments were conducted at MOIs of 0.01, 1 and 100. We observed a consistent pattern of evolution of bacteria and phage counts (Figure 3). The exponential growth of bacteria was followed by a rapid proliferation of phages. Then, the bacterial population collapsed between 6 and 10 h, while the phage population reached a plateau. An exception to this regular pattern was observed for experiments conducted with V1SA019 on USA300 at MOI of 0.01 where bacterial collapse was observed later, after about 18-20 h and slight decline of phage population was visible at 10-12 h. In several experiments, a regrowth of bacteria was observed, suggesting emergence of bacterial resistance. Regrowth rate appeared to increase with increasing MOI, which was confirmed by the modeling (see below). The final phage concentration was about 10^9^ - 10^11^ PFU.mL^-1^ depending on the phage/bacteria couple considered, irrespective of MOI. A slight decline of phage plateau was visible for V1SA019 on SH1000, especially for MOI of 100.

**Figure 3.**
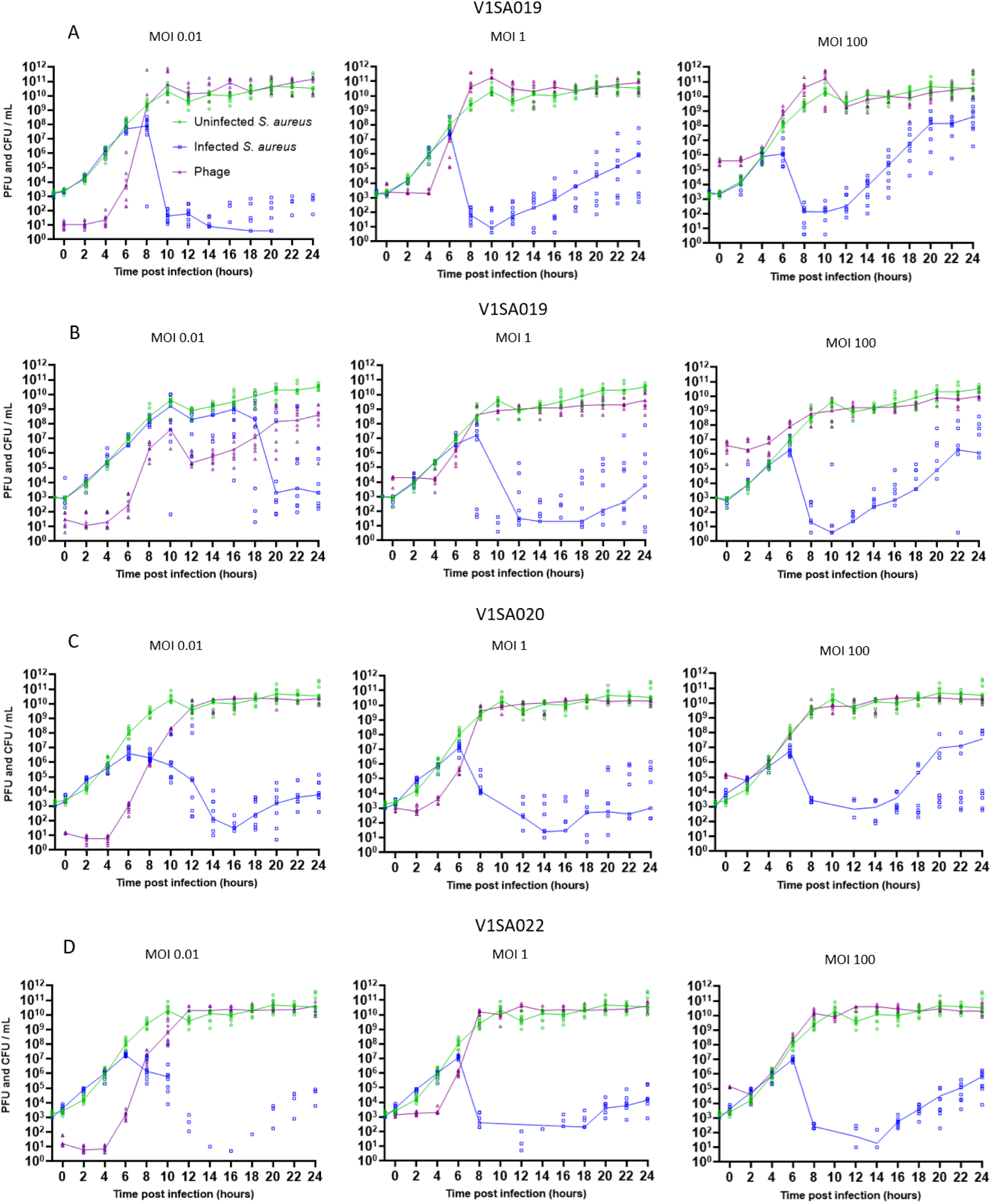
*in vitro* time-kill experiments between V1SA019 and SH1000(A) or USA300 (B), V1SA020 and SH1000 (C), V1SA022 and SH1000 (D). Each symbol represents an experimental point. Connecting line represents the median. Green: Enumeration of S. *aureus* in absence of phage (control condition). Blue: Enumeration of S. *aureus* in the presence of phage. Purple: Free phages in the presence of S. *aureus*.

### Population PK/PD modeling

The *S. aureus* SH1000 / phage V1SA019 pair was used as an illustration for the PK/PD modeling in the manuscript, while results for the other bacteria/phage pairs are provided as supplementary material.

The four-equation, eight-parameter model fitted individual bacteria and phages data very well, as shown in Figure 4 for the V1SA019 / SH1000 pair, and supplementary Figure S3 and S4 for the other pairs. Model parameters are shown in for the SH1000 / V1SA019 pair and summarized in supplementary Table S3 for all experiments. The phage production batch showed no influence on PK/PD parameters in all four models. There were minor differences in the final model between the four experiments, regarding Bmax parameter (fixed for V1SA019 / SH1000 Table 1 Line 2, random for the other pairs; Table S3 Line 2) and the error model (Table 1 Line 16, 17, 18; Table S3, Line 17, 18, 19). Overall, most parameters were estimated with acceptable precision (relative standard error < 50 %, Table 1), including inter-experiment variability. Large imprecision was observed for the median mutation rate (except for V1SA019 / SH1000 pairs, Table 1 Line 3; Table S3 Line 3). This was likely due to the high interexperiment variability of this parameter (high omega values, Table S3 Line 11). A common feature between the four models was a phage binding rate of resistant bacteria (bind_R_) lower than that for susceptible bacteria (bind_S_) (Table 1 and S3, Line 4 and 5), resulting in higher proliferation and inundation thresholds for resistant compared to susceptible bacteria (Table 1 Line 19 to 23; Table S3, Line 20 to 24). Also, the estimated typical burst size was larger than the values directly obtained from experimental data (Figure S2, Table S2,Table 1 and Table S3 Line 7). By comparing the results obtained for the SH1000 strain with the three phage candidates (V1SA019, V1SA020, V1SA022), differences between phages can be observed in terms of mutation rate, burst size and decay (Table S3 Line 3, 7, 8). The comparison of the V1SA019 / SH1000 and V1SA019 / USA300 model parameters also revealed differences associated with the bacterial strains. Compared to the SH1000 strain, the USA300 strain displayed a 10-fold lower estimated median burst size and much lower median mutation rate (Table S3, Line 7 and 3). However, considerable variability was observed for the mutation rate for the V1SA019 / USA300 experiments, with values ranging from 0 to 5.25 x 10^-8^ between experiments (median = 1.26 x 10^-13^) (Table S3, Line 3).

**Figure 4.**
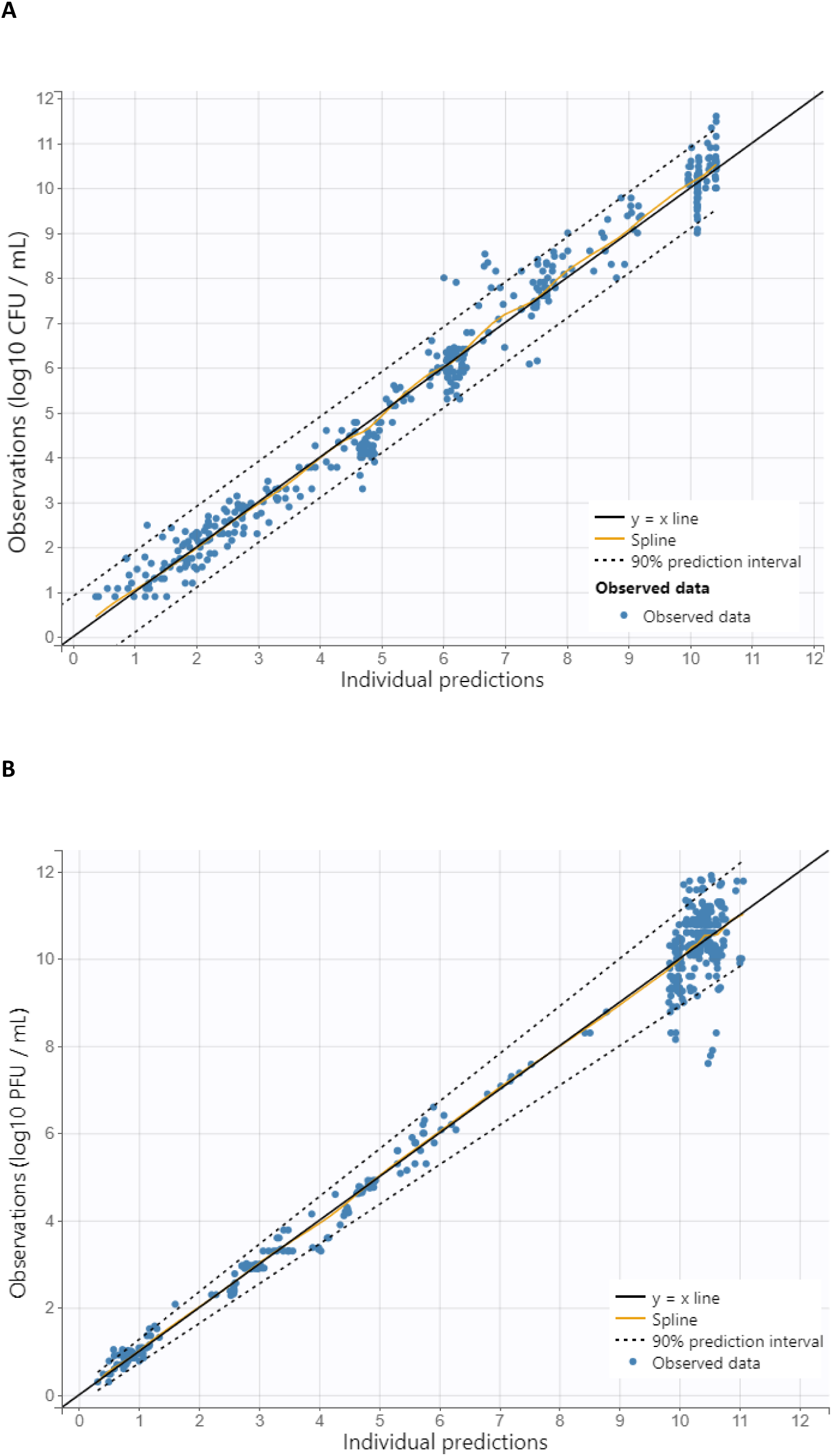
Observations versus model-based predictions of bacteria (A) and phage (B) counts for the V1SA019 / SH1000 pair. The blue dots indicate observation-prediction pairs. The black solid line is the y = x line, the yellow line is the spline regression line. The dashed lines indicate the 90 % prediction interval.

**Table 1.**
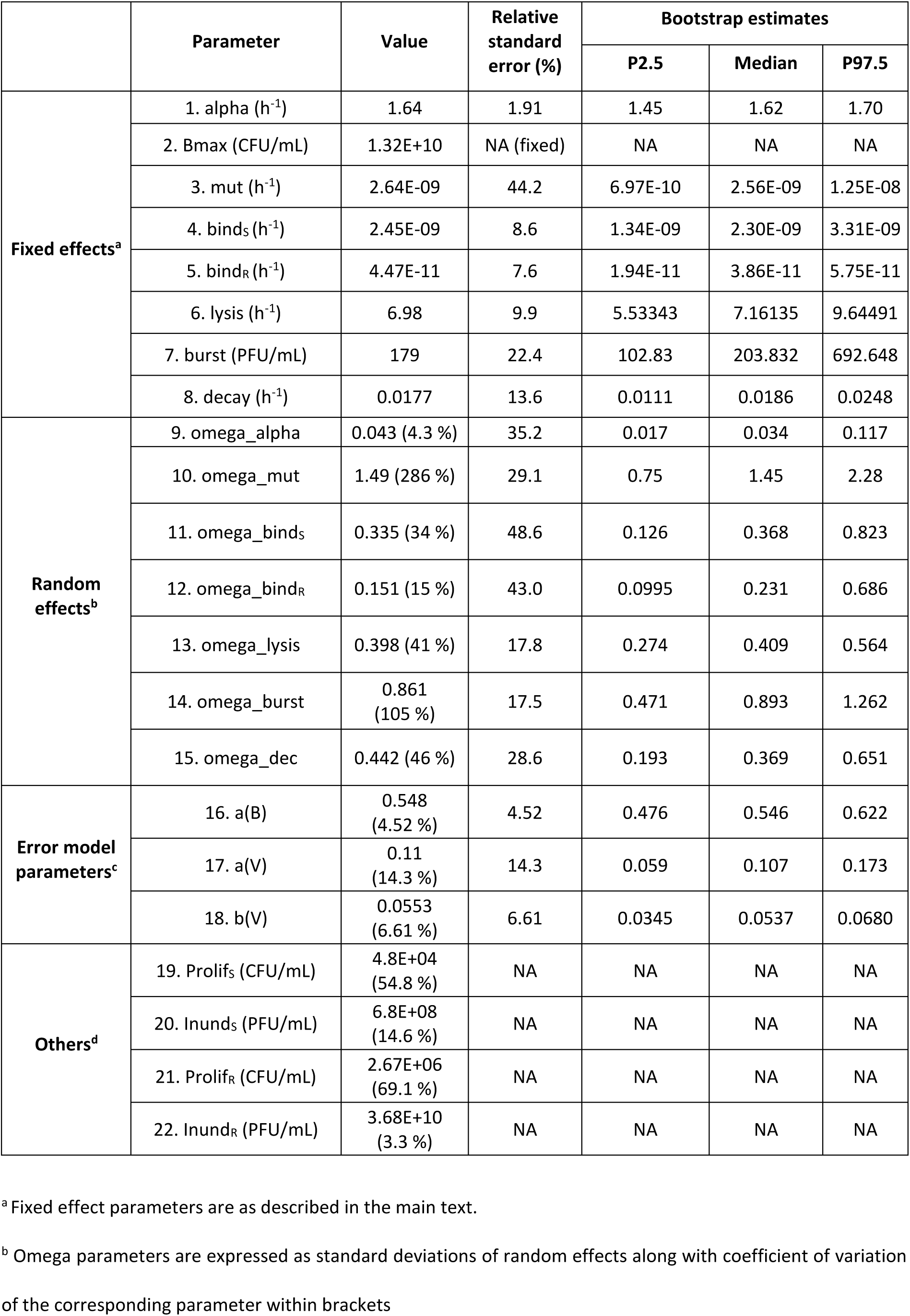
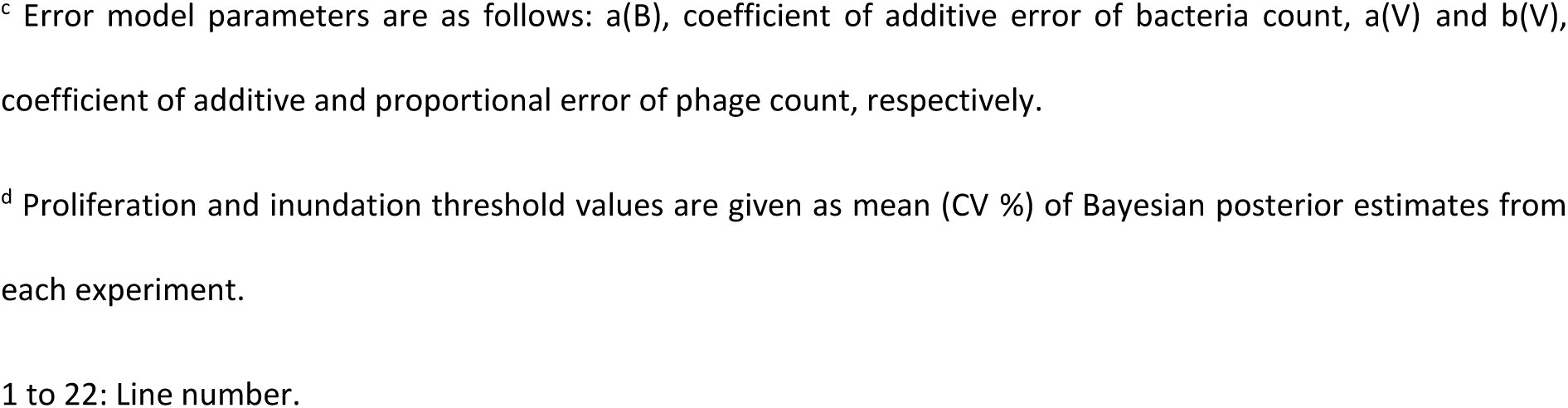
Parameter estimates of the phage-bacteria co-dynamics for *S. aureus* SH1000 / V1SA019.

|  | Parameter | Value | Relative standard error (%) | Bootstrap estimates |  |  |
| --- | --- | --- | --- | --- | --- | --- |
|  |  |  |  | P2.5 | Median | P97.5 |
| Fixed effects<sup>a</sup> | 1. alpha (h<sup>-1</sup>) | 1.64 | 1.91 | 1.45 | 1.62 | 1.70 |
|  | 2. Bmax (CFU/mL) | 1.32E+10 | NA (fixed) | NA | NA | NA |
|  | 3. mut (h<sup>-1</sup>) | 2.64E-09 | 44.2 | 6.97E-10 | 2.56E-09 | 1.25E-08 |
|  | 4. bind<sub>S</sub> (h<sup>-1</sup>) | 2.45E-09 | 8.6 | 1.34E-09 | 2.30E-09 | 3.31E-09 |
|  | 5. bind<sub>R</sub> (h<sup>-1</sup>) | 4.47E-11 | 7.6 | 1.94E-11 | 3.86E-11 | 5.75E-11 |
|  | 6. lysis (h<sup>-1</sup>) | 6.98 | 9.9 | 5.53343 | 7.16135 | 9.64491 |
|  | 7. burst (PFU/mL) | 179 | 22.4 | 102.83 | 203.832 | 692.648 |
|  | 8. decay (h<sup>-1</sup>) | 0.0177 | 13.6 | 0.0111 | 0.0186 | 0.0248 |
| Random effects<sup>b</sup> | 9. omega_alpha | 0.043 (4.3 %) | 35.2 | 0.017 | 0.034 | 0.117 |
|  | 10. omega_mut | 1.49 (286 %) | 29.1 | 0.75 | 1.45 | 2.28 |
|  | 11. omega_bind<sub>S</sub> | 0.335 (34 %) | 48.6 | 0.126 | 0.368 | 0.823 |
|  | 12. omega_bind<sub>R</sub> | 0.151 (15 %) | 43.0 | 0.0995 | 0.231 | 0.686 |
|  | 13. omega_lysis | 0.398 (41 %) | 17.8 | 0.274 | 0.409 | 0.564 |
|  | 14. omega_burst | 0.861 (105 %) | 17.5 | 0.471 | 0.893 | 1.262 |
|  | 15. omega_dec | 0.442 (46 %) | 28.6 | 0.193 | 0.369 | 0.651 |
| Error model parameters<sup>c</sup> | 16. a(B) | 0.548 (4.52 %) | 4.52 | 0.476 | 0.546 | 0.622 |
|  | 17. a(V) | 0.11 (14.3 %) | 14.3 | 0.059 | 0.107 | 0.173 |
|  | 18. b(V) | 0.0553 (6.61 %) | 6.61 | 0.0345 | 0.0537 | 0.0680 |
| Others<sup>d</sup> | 19. Prolif<sub>S</sub> (CFU/mL) | 4.8E+04 (54.8 %) | NA | NA | NA | NA |
|  | 20. Inund<sub>S</sub> (PFU/mL) | 6.8E+08 (14.6 %) | NA | NA | NA | NA |
|  | 21. Prolif<sub>R</sub> (CFU/mL) | 2.67E+06 (69.1 %) | NA | NA | NA | NA |
|  | 22. Inund<sub>R</sub> (PFU/mL) | 3.68E+10 (3.3 %) | NA | NA | NA | NA |

<sup>a</sup> Fixed effect parameters are as described in the main text.
<sup>b</sup> Omega parameters are expressed as standard deviations of random effects along with coefficient of variation
of the corresponding parameter within brackets
<sup>c</sup> Error model parameters are as follows: $a(B)$ , coefficient of additive error of bacteria count, $a(V)$ and $b(V)$ , coefficient of additive and proportional error of phage count, respectively.
<sup>d</sup> Proliferation and inundation threshold values are given as mean (CV %) of Bayesian posterior estimates from each experiment.
1 to 22: Line number.

Visual predictive checks shown in Figure **5** for V1SA019 / SH1000 supported internal validation of the model. Those plots also exhibited different patterns of bacterial dynamics, including a sharp decline followed by regrowth of total bacteria count after 10-12 h, with variable slopes. Another pattern of constant growth up to a maximum value corresponded to control experiments conducted with bacteria only (no phage).

**Figure 5.**
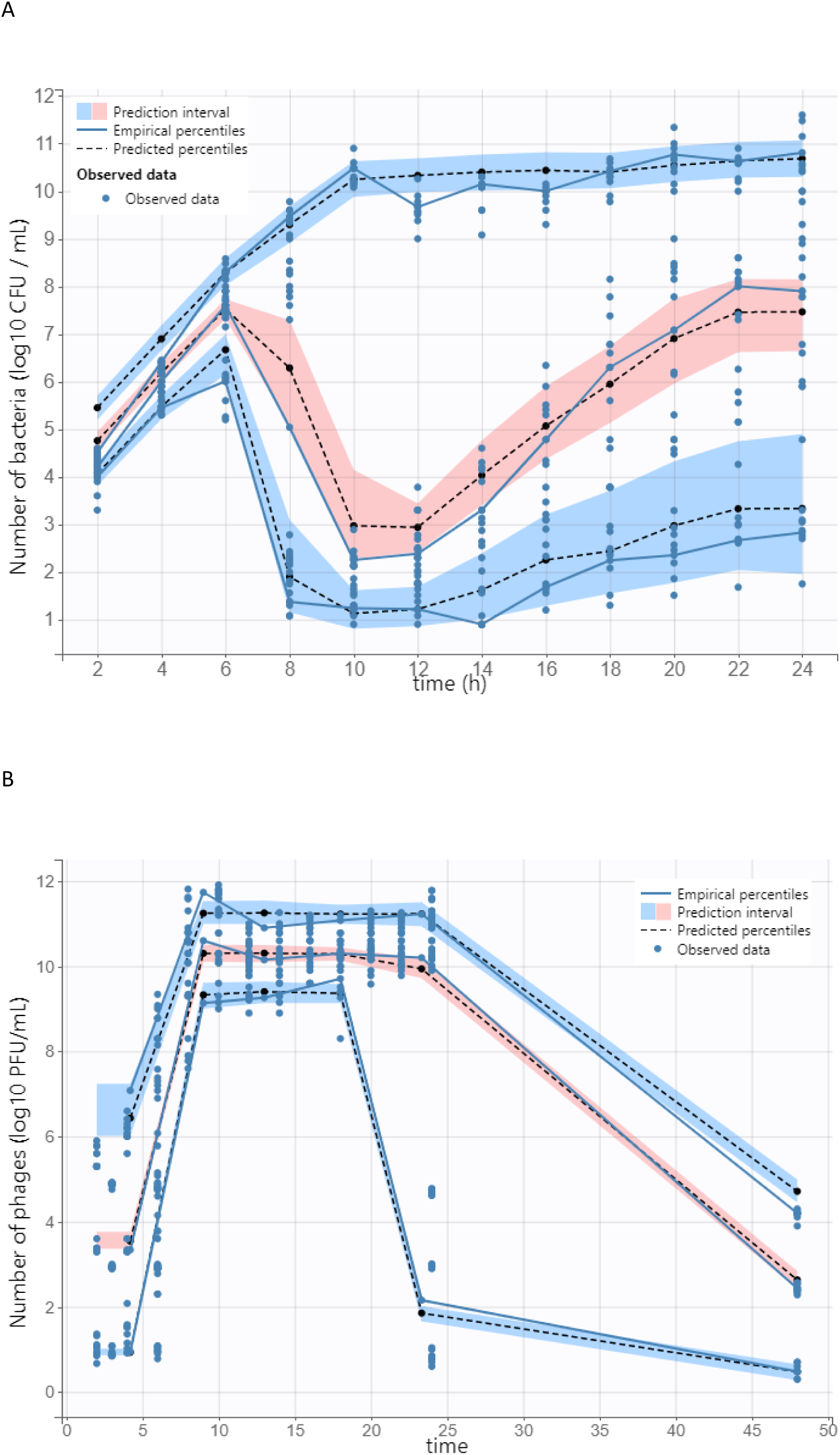
Visual predictive checks obtained with the final model for the V1SA019 / SH1000 pair. A and B panels show VPC for bacteria and phage, respectively. Blue dots are observed data. Blue solid and black dashed lines are 10^th^, 50^th^ (median) and 90^th^ percentiles of observed and simulated data (n = 1000 for each experiment), respectively. Blue and pink colored areas are 90 % prediction intervals of corresponding simulated percentiles.

The model also well described the time-course of the bacteria and phage counts of individual experiments. A selection of individual fits for various initial MOIs is shown in Figure S5 for our reference pair V1SA019 / SH1000. Those plots illustrate the bacterial regrowth that appeared to increase with increasing phage dose. This was confirmed by simulations performed with the final model parameters for various phage doses, shown in Figure 6. While regrowth was generally limited for an initial MOI of 0.01, it was dramatic for initial MOIs of 1 and 100.

**Figure 6.**
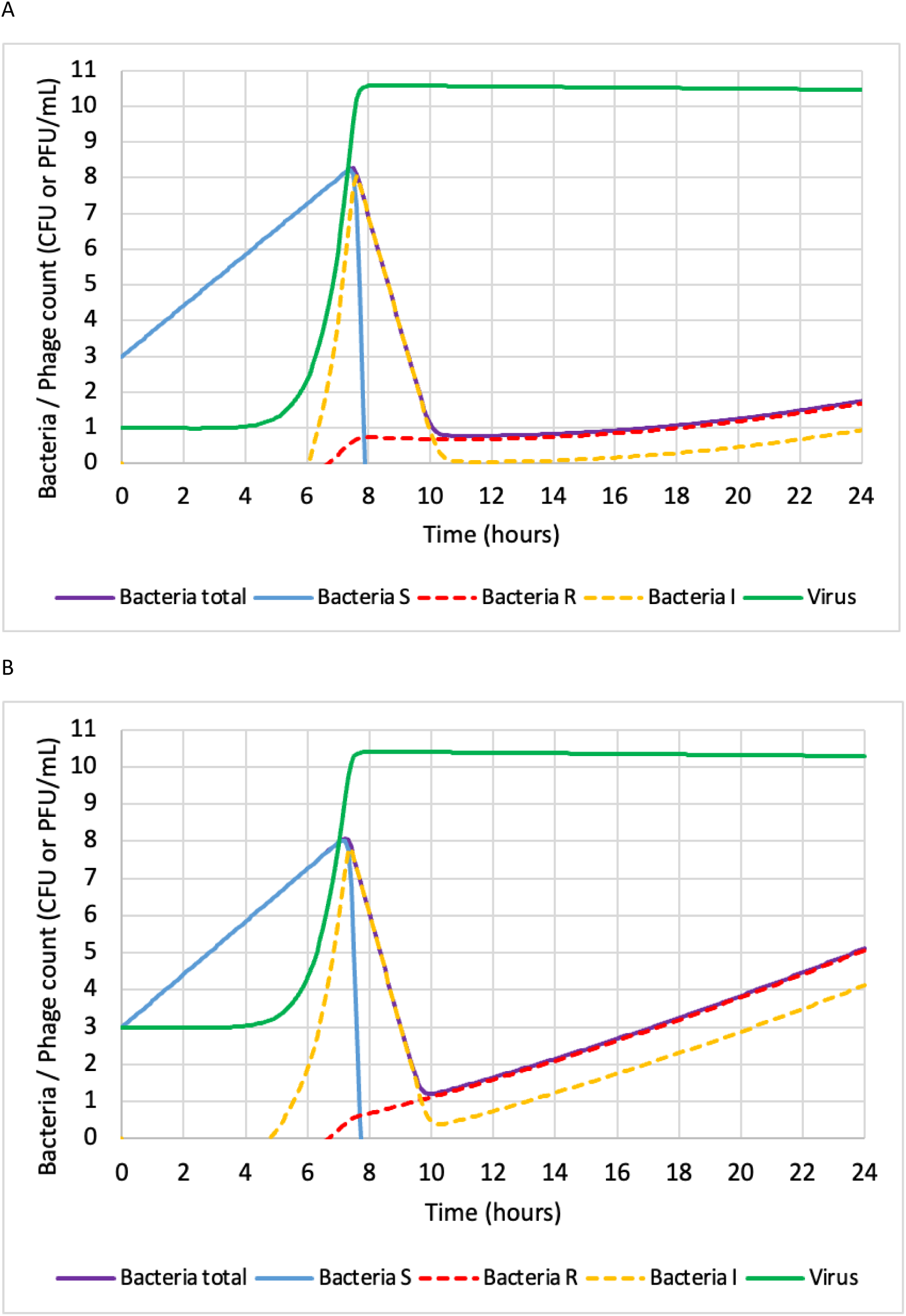
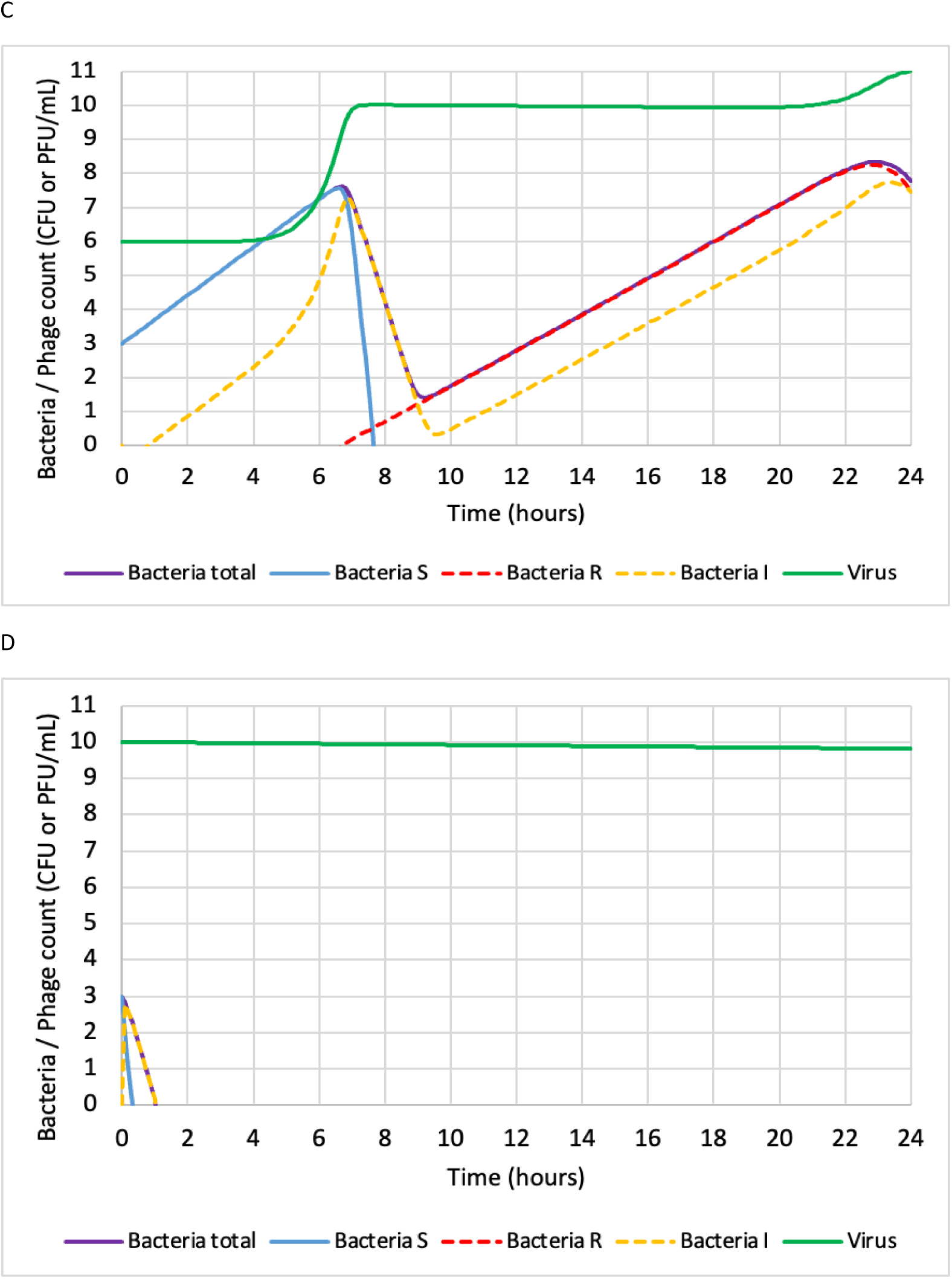
Simulated dynamics of bacteria and phage Simulations were performed with typical parameter values of the final model for V1SA019 / SH1000 reported in. Initial number of susceptible bacteria is 10^3^ in all simulations. Initial number of phages is 10, 10^3^, 10^6^ and 10^10^ in panel A, B, C, and D, respectively.

Bacterial regrowth rate appeared to be dependent on the initial phage concentration. Such regrowth was driven by the emergence of the resistant bacterial population to phages. However, a phage dose above the inundation threshold (10^10^ PFU mL^-1^) resulted in very fast killing of all bacteria and no regrowth according to simulations.

### Emergence of a resistant bacterial subpopulations

At the end of the experiments between V1SA019 and SH1000, one clone from each of the replicates of the three initial MOIs (i.e. three clones per MOI) were randomly selected, isolated, purified and sequenced. Resistance to V1SA019 was confirmed using the spot test assay with no lysis observed. Fitness of phage-resistant clones was assessed in comparison with the wild-type strain after overnight growth in caMH medium. Phage-resistant clones exhibited a delayed onset of exponential growth and reached a lower final OD_600_ compared to the wild-type (Figure S6). Each clone carried at least one mutation (non-synonymous or deletion) in one of the genes involved in the teichoic acid biosynthesis pathway, namely *tarA, tarK* or *tagO*, according to the KEGG pathway (KEGG pathway database) (Table S4).

The model predicted the appearance of resistant bacteria as early as 8 h after infection, irrespective of the initial MOI. After 8, 10, 12, 14 and 24 h of infection, up to 60 clones per time were isolated and tested for susceptibility to phage V1SA019 via the liquid phage susceptibility test. At 8 h, 0 %, 7 % and 7 % (60, 27 and 30 clones tested) resistant clones were identified for MOI of 0.01, 1 and 100 respectively. At 10 h, 0 %, 0 % and 19 % (60, 15 and 27 clones tested) resistant clones were identified for MOI of 0.01, 1 and 100 respectively. At 12h, 0 % resistant clones were identified for MOI of 0.01 and 1 (30 and 22 clones tested) but this percentage increased to 33 % and 100 % (3 and 10 clones tested) at 14 h for MOI 0.01 and 1 respectively. At 24 h, 96 % of resistant clones were identified for MOI of 0.01 and 100 % for MOI of 1 and 100 (24, 30 and 30 clones tested respectively) (Table S5).

## Discussion

### Co-dynamics of bacteria and phage

We combined *in vitro* experiments and mathematical modeling to investigate the co-dynamics between two *Staphylococcus aureus* strains and three candidate phages [10] across three MOIs. *In vitro* experiments revealed the emergence of a phage-resistant subpopulation across all phage–bacteria pairs tested. According to the mathematical model, this resistant subpopulation exhibited a reduced phage adsorption rate compared to susceptible bacteria, as well as distinct values for proliferation and inundation thresholds.

Our analysis confirmed that the co-dynamics between phages and bacteria can be effectively described using ODE, capturing predator–prey-like interactions. The model was inspired by the framework developed by Cairns and colleagues [5], and builds upon previous mathematical approaches established through *in silico* and *in vitro* studies of pathogens such as *Salmonella typhimurium* and *Campylobacter jejuni* [5], [12], [16], [17], [18], [19]. While probabilistic models—such as logistic regression approaches—have been applied to predict *S. aureus* inactivation by phages in matrices like pasteurized milk [20], to our knowledge, no ODE-based model has yet been developed to characterize the dynamic interactions between *S. aureus* and lytic phages under laboratory conditions. In addition, we employed a non-linear mixed-effects modeling approach to quantify experimental variability in pharmacokinetic/pharmacodynamic (PK/PD) parameters. This allowed us to capture substantial inter-experimental variability, particularly in mutation rate and phage burst size. It is worth noting that the estimated median mutation rate (2.64 × 10⁻⁹ / h) is consistent with previously reported *in vitro* mutation rates for *S. aureus* populations, which range from 1 × 10⁻⁸/h to 5 × 10⁻⁹ / h.

A key strength of our study lies in the simultaneous investigation of three distinct phage candidates and two *Staphylococcus aureus* strains. Our findings highlight notable differences in key dynamic parameters, particularly between the SH1000 and USA300 strains. These results suggest that phage–bacteria interaction dynamics are highly pair-specific and cannot be reliably generalized or extrapolated from a single strain–phage combination.

### Proliferation and inundation thresholds

The model enabled estimation of the inundation and proliferation thresholds that define two distinct modes of phage therapy—passive and active, respectively [4], [21]. In passive therapy, a sufficiently high initial phage dose ensures immediate bacterial clearance, while in active therapy, therapeutic success relies on *in situ* phage amplification following bacterial infection. *In vitro* time-kill assays showed that bacterial growth persisted in the presence of lytic phages until the inundation threshold was reached, at which point rapid bacterial killing occurred. Previous models estimated the minimum phage concentration required to inactivate *S. aureus* in milk at approximately 8 log₁₀ PFU mL⁻¹ [20], a value closely matching the inundation thresholds calculated in the present study (8.0 to 8.7 log₁₀ PFU mL⁻¹), despite differing experimental conditions. In contrast, the proliferation threshold was more difficult to discern from time-course data, as previously noted by Cairns *et al*. [5], particularly since only free phages were quantified. Despite this limitation, our study relied on free phage measurements to estimate the proliferation threshold, which for susceptible bacteria ranged from 4.5 to 5.2 log₁₀ CFU mL⁻¹ and appeared independent of the phage, the initial phage dose, or the bacterial strain.

Although these thresholds may vary under different *in vitro* and *in vivo* conditions, this study demonstrates that in the context of active phage therapy, a minimal bacterial population is necessary to enable phage amplification, reach the inundation threshold, and ultimately reduce bacterial load [22]. This might suggest that different phage doses below the inundation threshold would produce similar bacterial load dynamics; however, this assumption is incorrect. Our results show a clear effect of the initial phage dose—still below the inundation threshold—on both the average final abundance of resistant bacteria and their growth rate. This phenomenon was observed consistently across all phage–bacteria pairs studied. Under certain conditions (e.g., phage V1SA019 infecting SH1000 at a MOI of 0.01), complete absence of resistant bacteria could be observed at the end of the experiment. This effect may result from the use of a closed culture system. Delayed control of the susceptible bacterial population at lower MOI leads to greater depletion of the culture medium, which was not replenished during our experiments, due to extensive and prolonged bacterial replication. Consequently, resistant bacteria emerging at low MOI encounter a less favorable environment for growth.

The correlation between bacteria regrowth and initial MOI is intriguing. Our simulations performed with various MOIs provided interesting insights (Figure 6) and suggested the importance of achieving the inundation threshold of the resistant subpopulation for controlling the regrowth with active phage therapy. For the lowest MOI (0.01), it took longer for the phage population to achieve the inundation threshold of susceptible bacteria. This leaves more time for susceptible bacteria to grow, and the susceptible bacteria peak is the highest, compared with higher MOI. The phage population plateau is also higher, close to the inundation threshold of resistant bacteria estimated at 4 x 10^10^ PFU ml^-1^ (median). By contrast, higher MOI result in lower susceptible bacteria peak and lower phage plateau, below the inundation threshold of resistant bacteria. Based on those observations, we assume that two ways exist to prevent the emergence of resistant bacteria and prevent bacteria regrowth in such *in vitro* conditions. The first one is to use a very high dose of phage, greater that the inundation threshold of susceptible bacteria that totally suppress the mutation process (passive therapy). The other option is to use active therapy with a phage dose that permits phage amplification up to the inundation threshold of resistant bacteria.

For phage doses exceeding the inundation threshold (passive therapy), bacterial concentrations are rapidly reduced due to lysis of the majority of susceptible cells. Although our simulation results indicate that administration of a phage dose above this threshold is necessary for rapid bacterial clearance without the emergence of resistance, reaching such high phage concentrations may be challenging due to formulation constraints and accessibility at the infection site. Furthermore, passive therapy may not achieve complete eradication, leaving a residual susceptible bacterial population with a reduced probability of encountering phages [23], [24]. Consequently, phage concentrations at the infection site may decline, allowing bacterial regrowth until the proliferation threshold is reached, thus enabling active therapy if viable phages remain. Obeso *et al*. reported that the efficacy of phage-mediated *S. aureus* inactivation in milk is influenced by the initial bacterial load, likely related to the bacterial proliferation threshold [20]. In contrast, Bigwood *et al.* demonstrated that biocontrol of low levels of *S. typhimurium* in liquid foods is achievable by applying sufficiently high phage concentrations, independent of initial pathogen counts [19]. Therefore, optimal phage dosing must balance immediate bactericidal efficacy with long-term control outcomes.

### Emergence of a resistant bacterial subpopulation

All nine clones isolated 24 h post-infection harbored mutations in genes involved in the teichoic acid biosynthesis pathway, which encodes the primary receptor for phages in *S. aureus* and, more generally, in Gram-positive bacteria [25], [26]. This finding indicates that, despite variations in bacterial population control across MOI conditions, phage-induced selective pressure consistently drives a similar adaptive response. Nonetheless, all resistant clones exhibited a growth defect when tested individually in caMH medium, suggesting a fitness cost that may facilitate their elimination by the host immune system, as previously observed in *in vivo* models of *S. aureus* under phage predation [27], [28]. Clones isolated at intermediate time points during the kinetics were not sequenced, and the presence of mixed subpopulations during these phases cannot be ruled out. Liquid susceptibility tests using V1SA019 on clones sampled at 8, 10, 12, and 14 h post-infection revealed a progressive shift from susceptible to resistant phenotypes, supporting the model’s prediction that the resistant subpopulation emerged and start to proliferate around 8 hours after infection onset.

Additional evidence supporting the presence of coexisting subpopulations during the kinetics comes from the observation that, in certain phage–bacterium pairs (e.g., V1SA019 and SH1000), the phage concentration does not simply reach a peak and remain stable, but instead shows a slight decline after 10 hours (by 0.5 to 1 log₁₀). This pattern, previously reported by Cairns *et al.* in a different phage–bacterium system [5], suggests a potentially generalizable phenomenon rather than an experimental artifact. One possible explanation is that phages may acquire mutations enabling them to infect the resistant bacterial population [29], [30]. An alternative explanation could involve the coexistence of phages with both susceptible and resistant bacterial subpopulations over time [31], [32]. These findings underscore the need for future studies to monitor the proportions of susceptible and resistant bacteria throughout the experiment and to perform sequencing of both bacterial and phage populations to detect the possible emergence of mutant lineages.

## Conclusion

Our mechanistic PK/PD model reliably captures the complex dynamics of phage – *S. aureus* interactions, including resistance emergence. Its relevance now warrants evaluation using *in vivo* data to support translational applicability.

## Acknowledgments

The authors acknowledge the scientific review service of Hospices Civils de Lyon for the English writing proofreading.

This study was supported by the French Agence Nationale pour la Recherche (PHAG-ONE project, ANR 20-PAMR-0009).

## Supplementary Materials and Methods

### Adsorption time assay and One-Step Growth Curve experiment (burst size)

Adsorption time is the time required for bacteriophage to be absorbed into host bacterial cells as first stage of phage infection. Adsorption time was set as the time required for 95% of the phage population to be attached to a bacterium. Phage adsorption tests were performed as described by Marashi and colleagues with few modifications [33]. Briefly, phages were added to late log phase cultures of SH1000 or USA300 strains in 25 mL of caMH in a 50 mL conical tubes (approximately 10^8^ CFU mL^-1^) at an approximate MOI of 0.01 and incubated at 37°C without agitation. Samples were taken every 4 min up to 45 min after phage addition and were passed through a 0.2 µm filter (Merk, Darmstadt, Germany) to remove *S. aureus* cells and bound phages. Unbound ‘free’ phages were counted using spot tests and double layer plaque assays. To estimate the latent period (which is defined as the time between absorption and the beginning of the first burst) and burst size (which is defined as the number of new phages produced and released by an infected bacterium) of a single round of phage replication, the same protocol was applied with modifications: culture was serial diluted in pre-warmed caMH at 1/100, 1/1000 and 1/10000 at “T= Adsorption time” according to Kropinski and colleagues [34]. Samples were taken every 6 min up to 2 h post phage addition. The data obtained from the most diluted flask were used during analyses to avoid re-adsorption phenomena. The burst size value was calculated as follows: “Average phage concentration at plateau/Number of adsorbed phages at T“ Adsorption time” [34]. The one-step growth curve experiment was carried out in triplicates using three phage production batches.

---

**Supplementary Figure S1.**
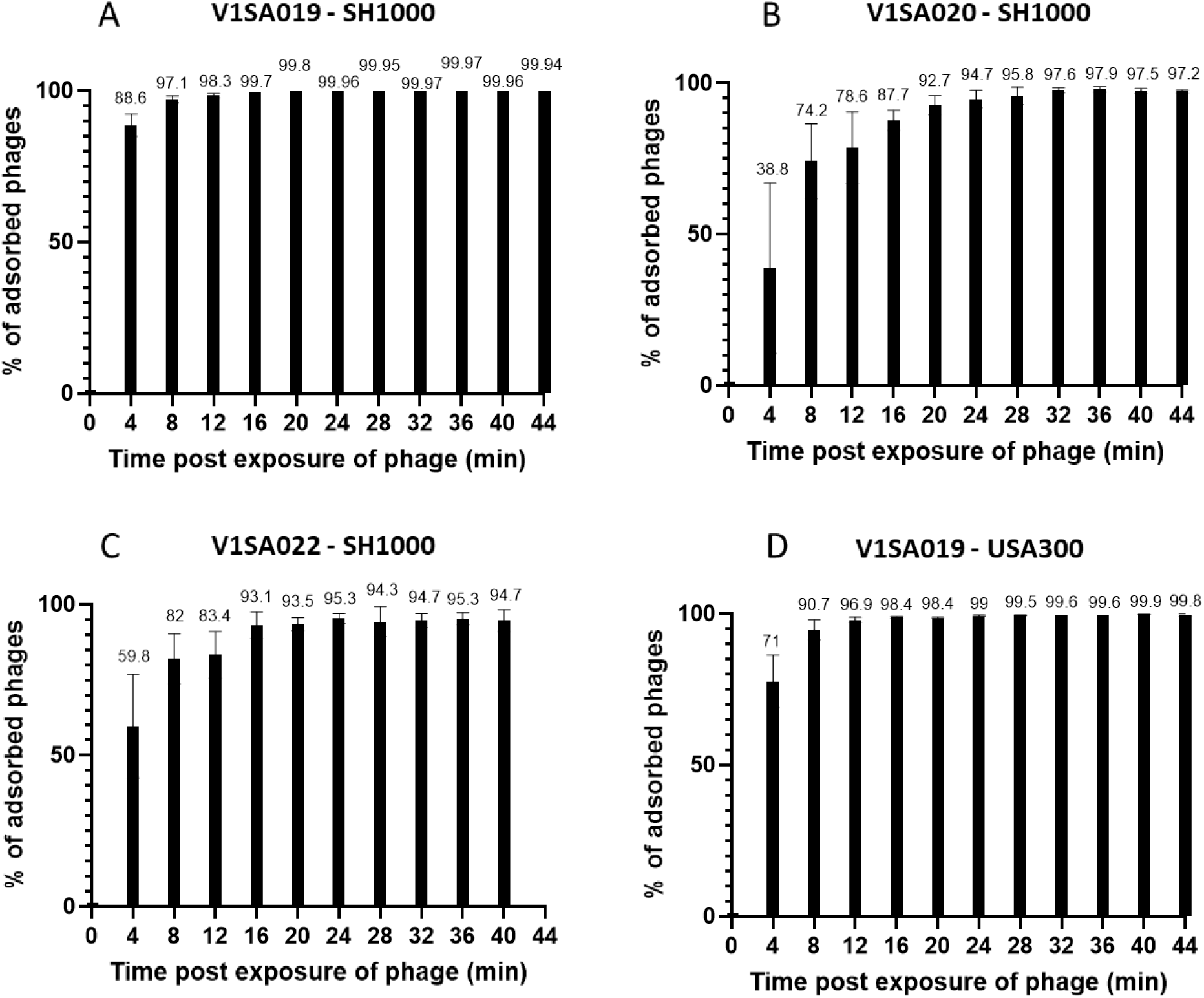
Adsorption kinetics for V1SA019 (A), V1SA020 (B), V1SA022 (C) phages on SH1000 strain and V1SA019 (D) on USA300 strain. Each graph represents the average of 3 independent experiments with standard deviation. The percentage of phage adsorption is indicated at the top of each column.

**Supplementary Figure S2.**
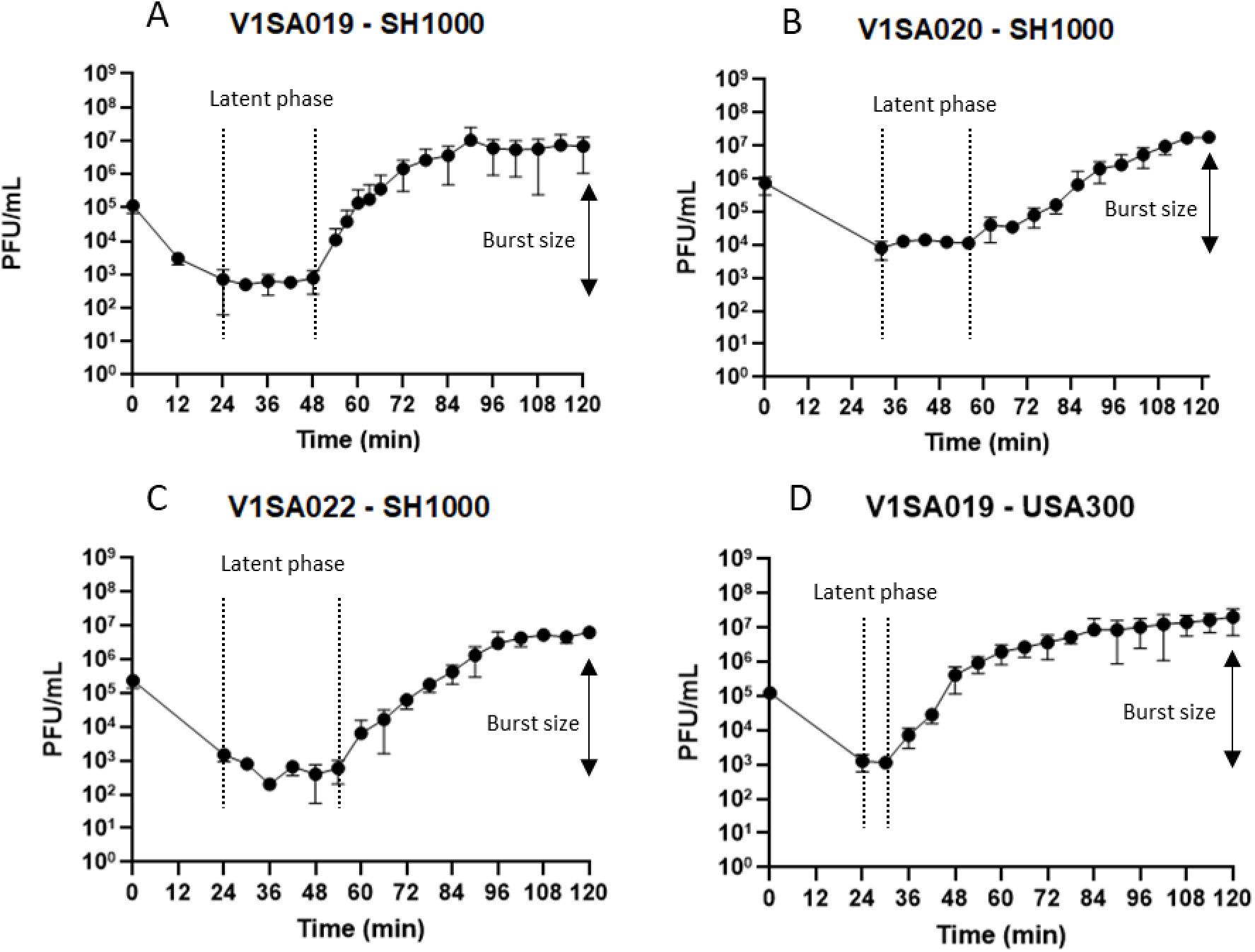
One-Step Growth Curve (for burst size value calculation) for V1SA019 (A), V1SA020 (B), V1SA022 (C) phages on SH1000 strain and V1SA019 (D) on USA300 strain. Each circle represents the average of 3 independent experiments with standard deviation. The latent phase is indicated between the dotted lines. The solid arrow represents the burst size.

**Supplementary figure S3.**
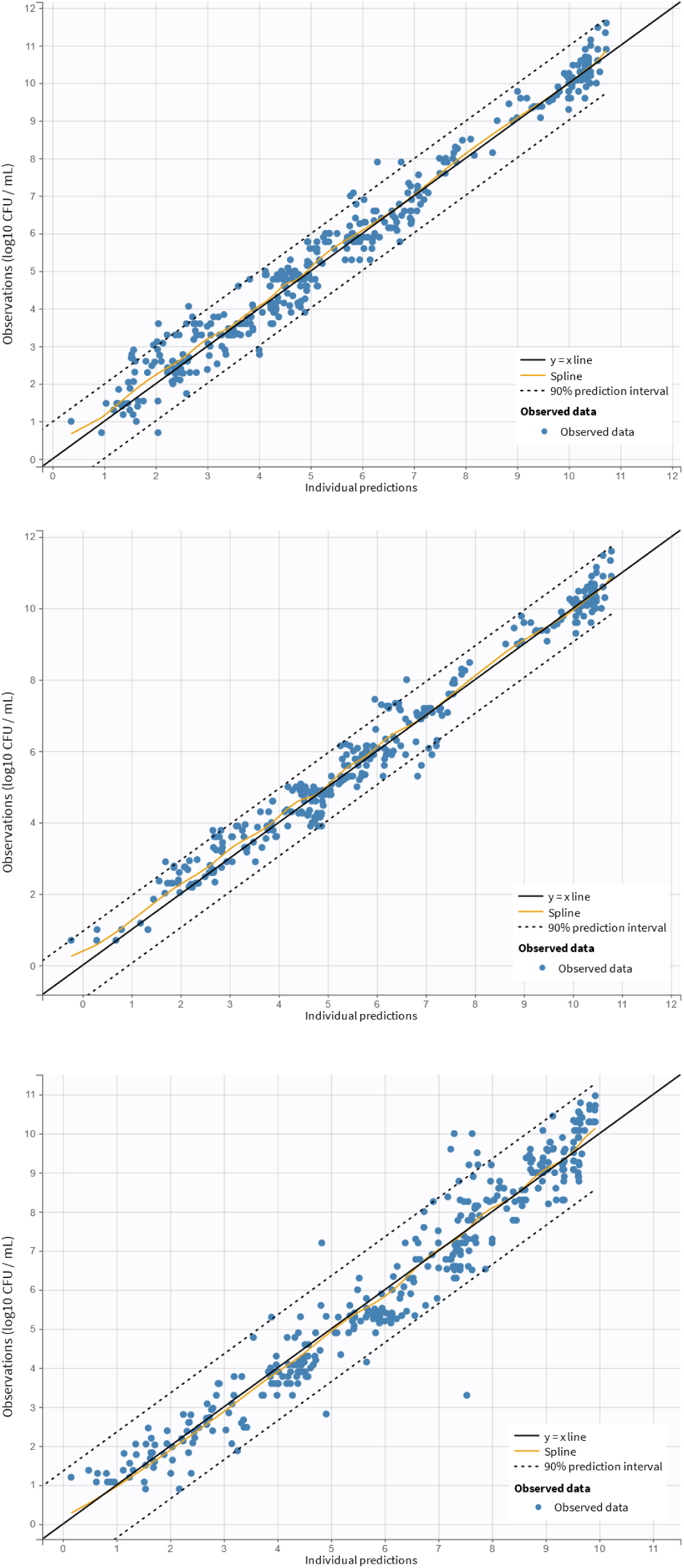
Observations versus model predictions of bacteria count Top, middle and bottom panel are bacteria counts for V1SA020 / SH1000, V1SA022 / SH1000, and V1SA019 / USA300 pairs, respectively.

**Supplementary figure S4.**
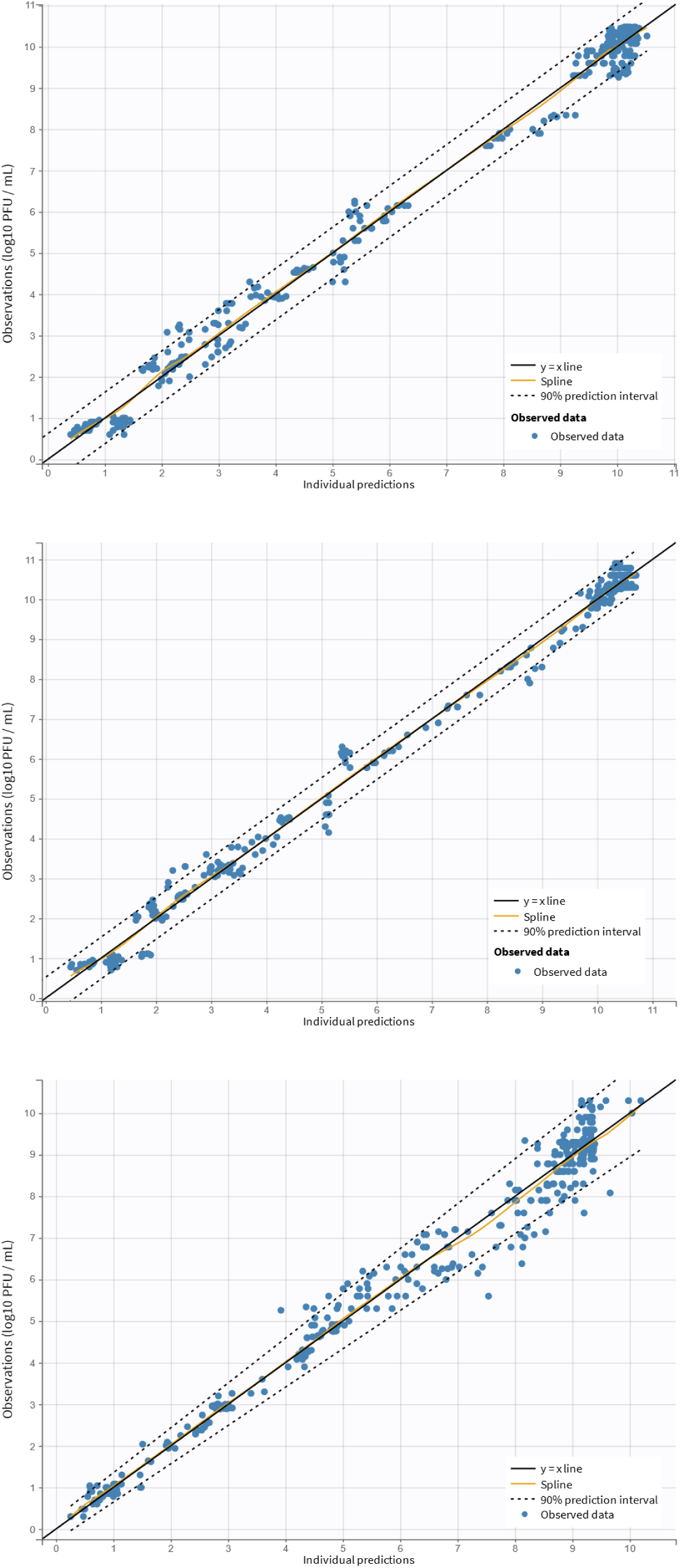
Observations versus model predictions of phage count Top, middle and bottom panel are phage counts for V1SA020 / SH1000, V1SA022 / SH1000, and V1SA019 / USA300 pairs, respectively.

**Supplementary Figure S5.**
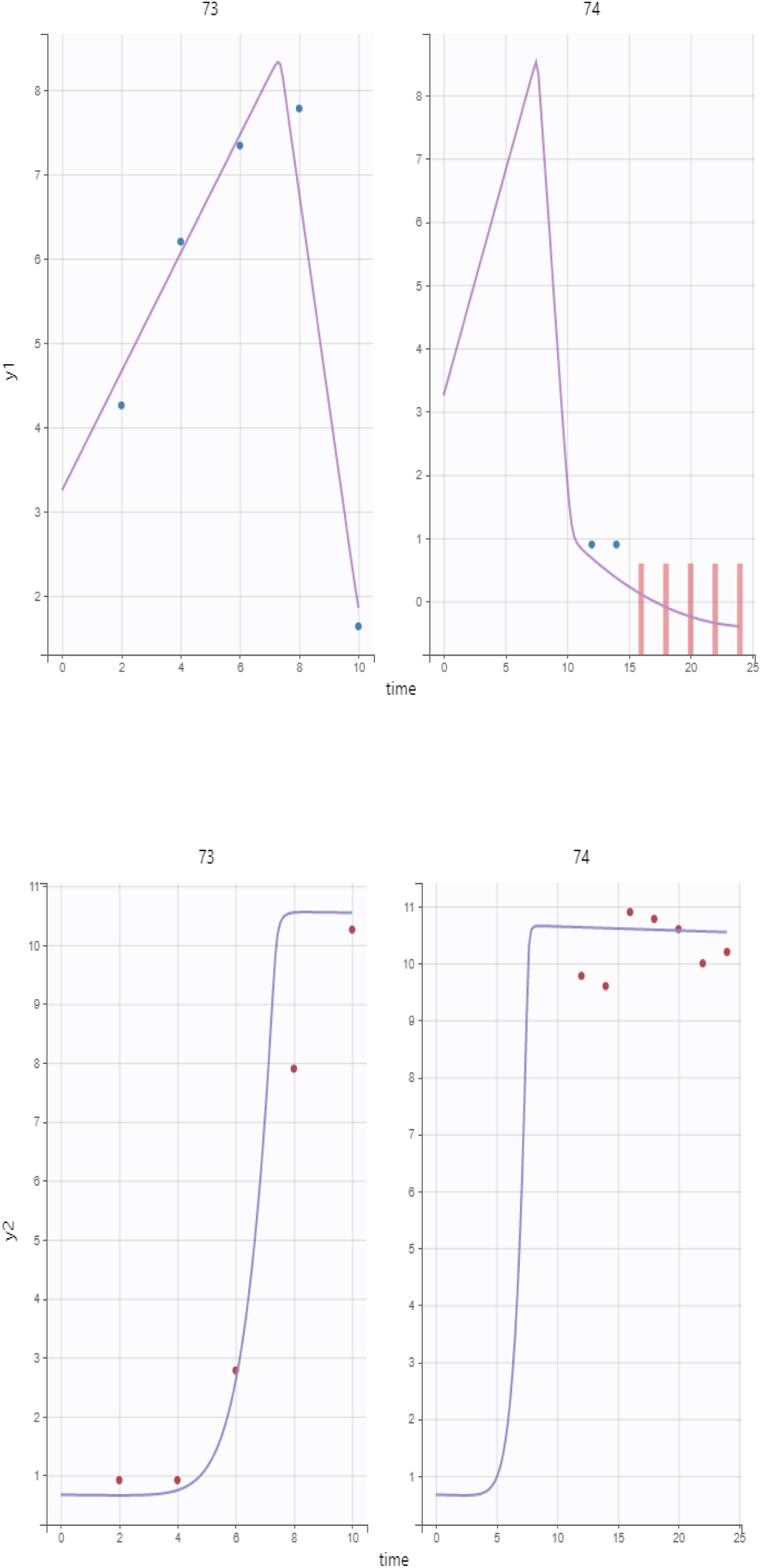
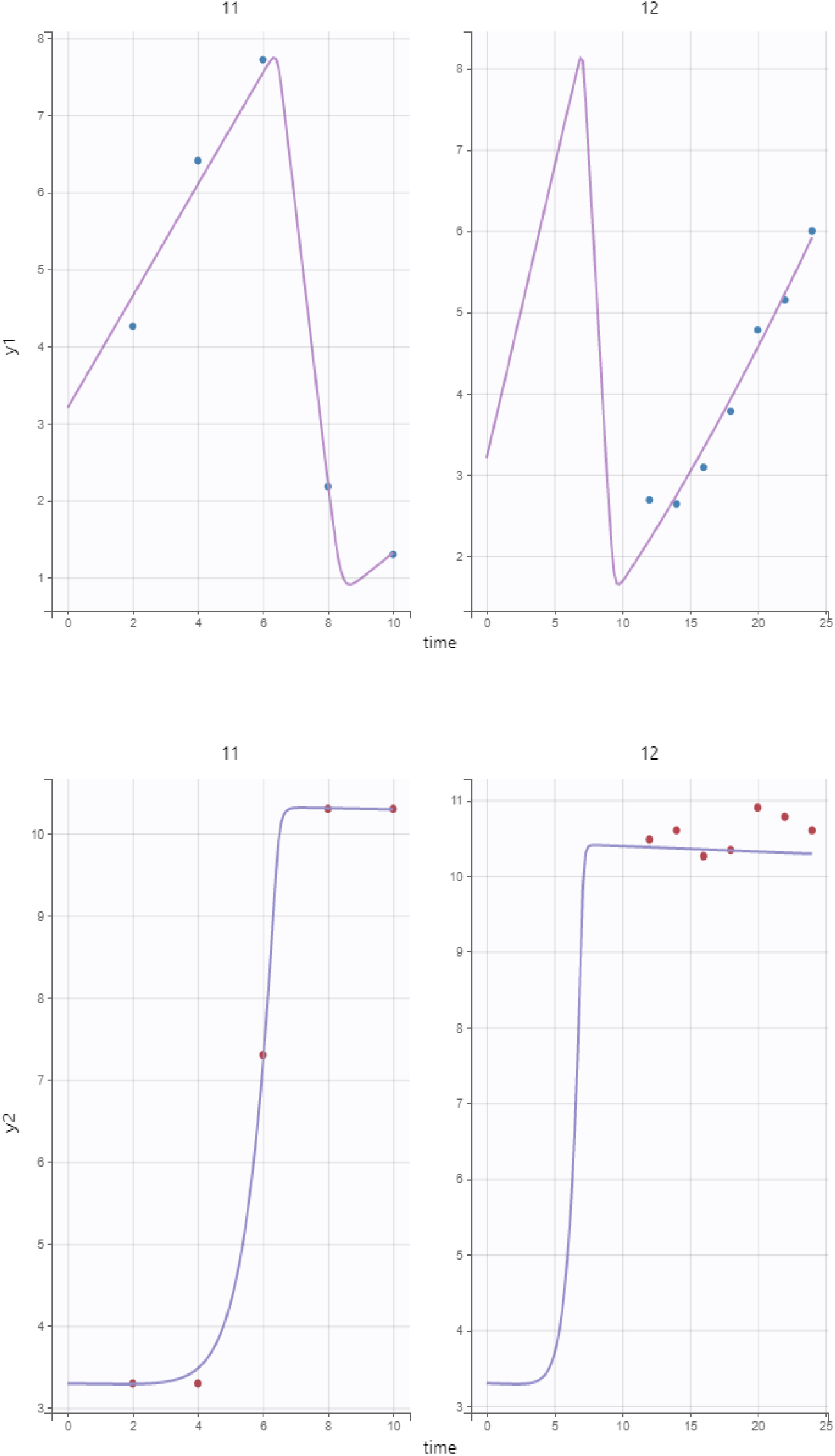
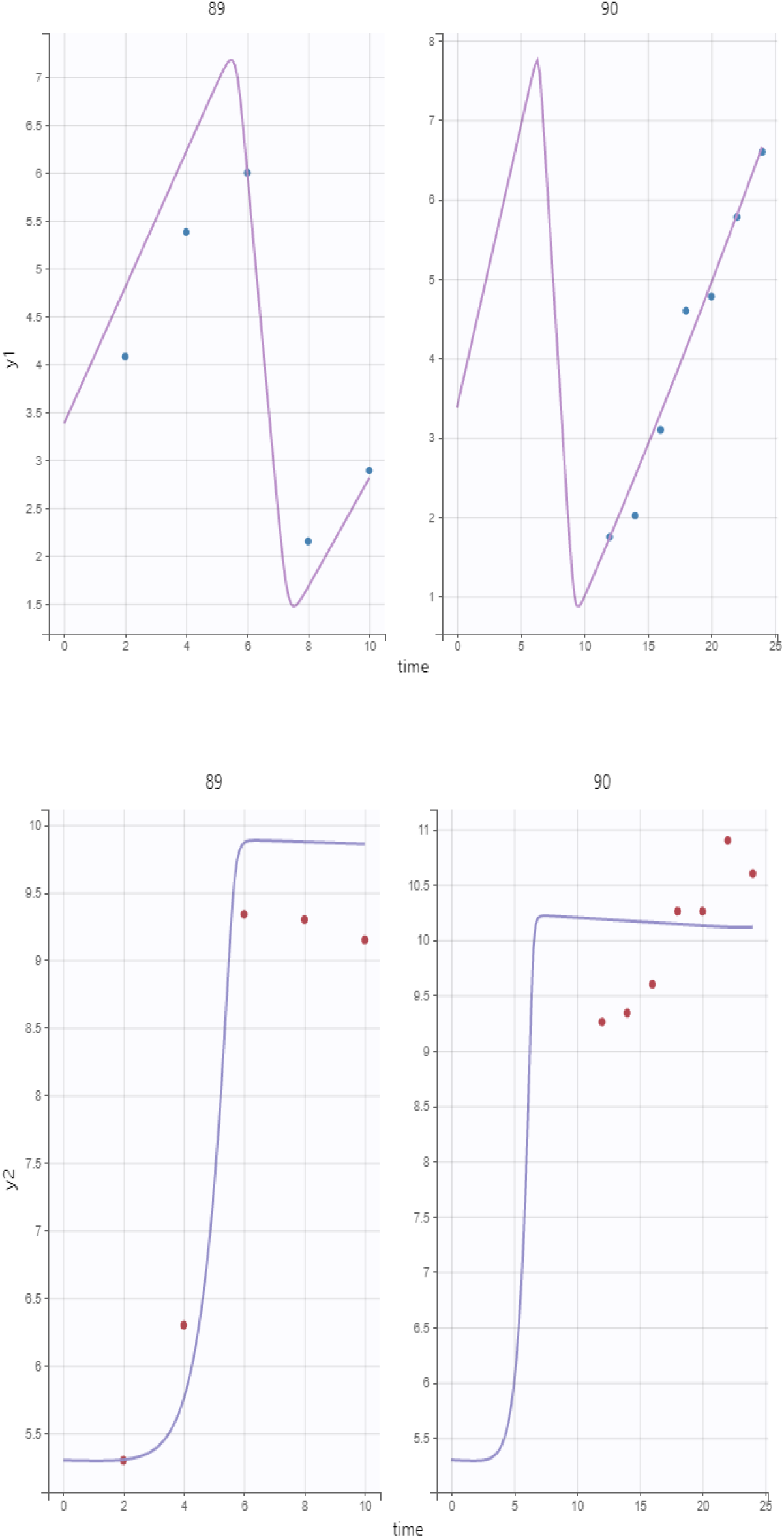
Examples of individual fits of bacteria and phage counts for the V1SA019 / SH1000 model Blue and red dots are bacteria (y1) and phage (y2) counts, respectively. Pink and purple lines are individual model predictions for bacteria and phage, respectively. Red bars indicate observations below the quantification limit. #73 and 74: B(0) = 1.8E03; V(0) = 5.0E00; MOI initial ≈ 0.01 #11 and 12: B(0) = 1.6E03; V(0) = 2.0E03; MOI initial ≈ 1 #89 and 90: B(0) = 2.4E03; V(0) = 2.0E05; MOI initial ≈ 100 Each pair represents two successive experiments with the same initial conditions and different time windows, 1-10 h and 12-24 h.

**Supplementary Figure S6.**
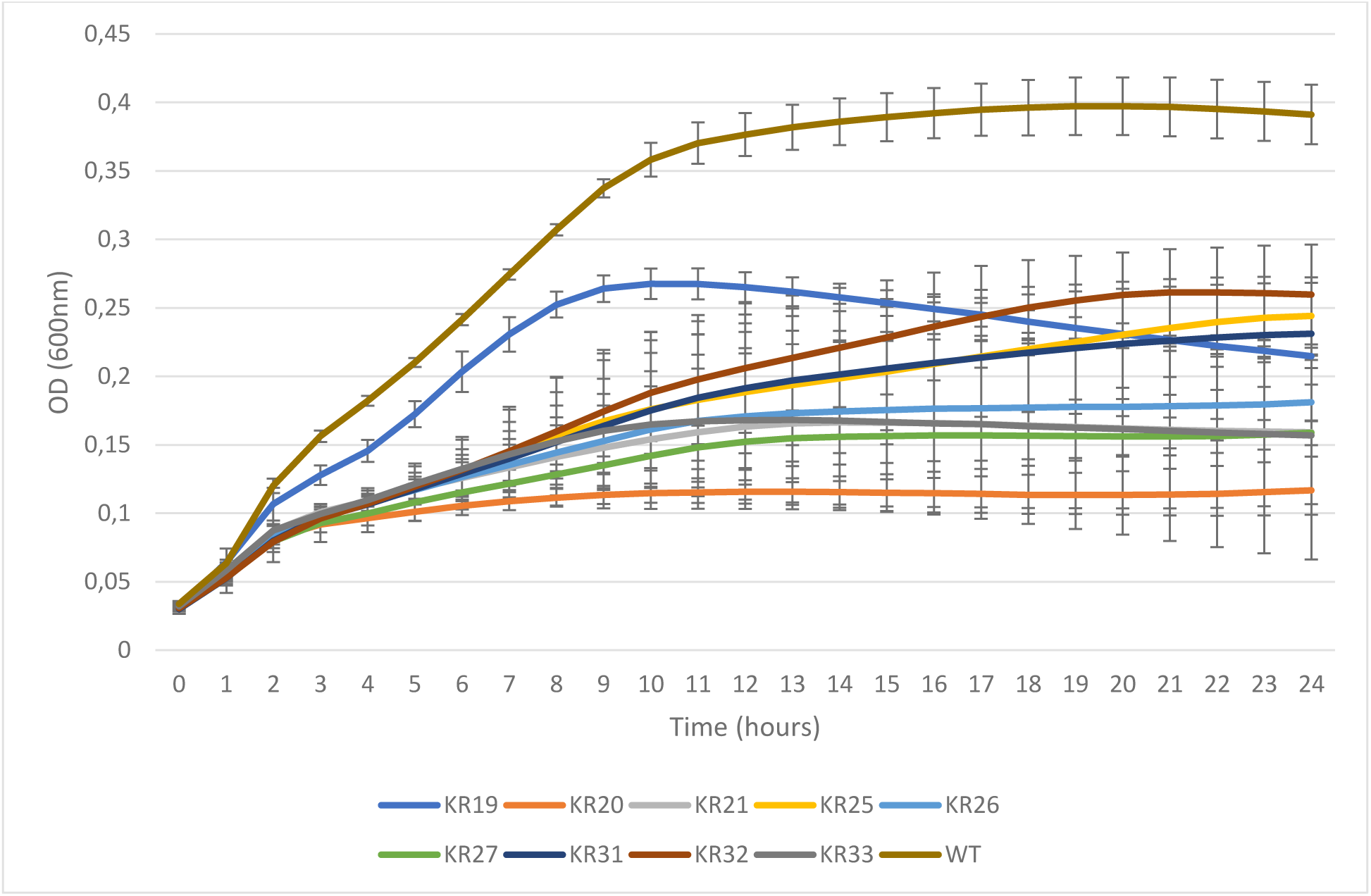
Growth curves for SH1000 resistant clones to V1SA019. Bars represent the standard deviation of 4 biological replicates. WT: Wild-type SH1000 strain.

**Supplementary Table S1.** Adsorption time. determined for phages V1SA019, V1SA020, V1SA022 on strains SH1000 and USA300. nd: not determined. Adsorption time is defined as the time where 95% of phages are adsorbed on bacteria.

| Adsorption time (min) |  | Bacterial strains |  |
| --- | --- | --- | --- |
|  |  | SH1000 | USA300 |
| Phages | V1SA019 | 8 | 12 |
|  | V1SA020 | 28 | nd |
|  | V1SA022 | 24 | nd |

**Supplementary Table S2.** Burst size values. determined for phages V1SA019, V1SA020, V1SA022 on strains SH1000 and USA300. Calculation based on data from One-Step Growth Curve experiment. nd: not determined.

| Burst size values |  | Bacterial strains |  |
| --- | --- | --- | --- |
|  |  | SH1000 | USA300 |
| Phages | V1SA019 | 42 | 134 |
|  | V1SA020 | 24 | nd |
|  | V1SA022 | 27 | nd |

**Supplementary Table S3.** Comparison of parameters of the phage-bacteria co-dynamics for the 4 bacteria/phage pairs.

|  | Parameter | SH1000 / V1SA019 | SH1000 / V1SA020 | SH1000 / V1SA022 | USA300 / V1SA019 |
| --- | --- | --- | --- | --- | --- |
| Fixed effects<sup>a</sup> | 1. alpha (h<sup>-1</sup>) | 1.64<br>(1.91 %) | 1.33<br>(3.92 %) | 1.31<br>(3.29 %) | 1.8<br>(2.66 %) |
|  | 2. Bmax (CFU/mL) | 1.32E+10<br>(NA, fixed) | 1.9E+10<br>(43.7 %) | 2.72E+10<br>(30.9 %) | 5.23e+08<br>(32.5 %) |
|  | 3. mut (h<sup>-1</sup>) | 2.64E-09<br>(44.2 %) | 1.46E-10<br>(382 %) | 6.85E-13<br>(>200 %) | 1.26e-13<br>(>200 %) |
|  | 4. binds (h<sup>-1</sup>) | 2.45E-09<br>(8.6 %) | 6.19e-09<br>(22 %) | 2.74E-09<br>(21.2 %) | 1.7e-08<br>(12.6 %) |
|  | 5. bind<sub>R</sub> (h<sup>-1</sup>) | 4.47E-11<br>(7.6 %) | 5.65e-11<br>(14.6 %) | 1.09E-11<br>(42.4 %) | 4.82e-10<br>(33.5 %) |
|  | 6. lysis (h<sup>-1</sup>) | 6.98<br>59.9 %) | 4.37<br>(8.13 %) | 5.96<br>(13.6 %) | 6.29<br>(12.7 %) |
|  | 7. burst (PFU/mL) | 179<br>(22.4 %) | 139<br>(45.6 %) | 503<br>(35 %) | 17.9<br>(33.9 %) |
|  | 8. decay (h<sup>-1</sup>) | 0.0177<br>(13.6 %) | 0.0336<br>(18.6 %) | 0.0499<br>(16.7 %) | 0.0109<br>(45.7 %) |
| Random effects<sup>b</sup> | 9. omega_alpha | 0.043<br>(35.2 %) | 0.207<br>(11.8 %) | 0.172<br>(11.6 %) | 0.108<br>(24.8 %) |
|  | 10. Omega_Bmax | NA | 0.82<br>(40.2 %) | 0.754<br>(32 %) | 1.63<br>(14 %) |

|  | Parameter | SH1000 /<br>V1SA019 | SH1000 /<br>V1SA020 | SH1000 /<br>V1SA022 | USA300 /<br>V1SA019 |
| --- | --- | --- | --- | --- | --- |
|  | 11. omega_mut | 1.49<br>(29.1 %) | 2.57<br>(46.8 %) | 4.59<br>(28.4 %) | 7.62<br>(29.7 %) |
|  | 12. omega_bind<sub>s</sub> | 0.335<br>(48.6 %) | 1.17<br>(15.8 %) | 1.35<br>(14.5 %) | 0.604<br>(18.4 %) |
|  | 13. omega_bind<sub>R</sub> | 0.151<br>(43.0 %) | 0.243<br>(27.9 %) | 0.745<br>(38.2 %) | 0.962<br>(33.1 %) |
|  | 14. omega_lysis | 0.398<br>(17.8 %) | 0.427<br>(15.1 %) | 0.634<br>(17.6 %) | 0.599<br>(27 %) |
|  | 15. omega_burst | 0.861<br>(17.5 %) | 0.963<br>(17.1 %) | 1.25<br>(21.9 %) | 1.48<br>(13.8 %) |
|  | 16. omega_dec | 0.442<br>(28.6 %) | 0.53<br>(29.1 %) | 0.492<br>(33 %) | 1.41<br>(21.5 %) |
| **Error model**<br>**parameters<sup>c</sup>** | 17. a(B) | 0.548 | 0.6 | 0.575 | 0.824 |
|  | 18. a(V) | 0.11 | 0.38 | 0.319 | 0.168 |
|  | 19. b(V) | 0.0553 | NA | NA | 0.0475 |
| **Others<sup>d</sup>** | 20. Prolif<sub>s</sub><br>(CFU/mL) | 4.8E+04<br>(54.8 %) | 7.00E+04<br>(174.9 %) | 4.78E+04<br>(63.8 %) | 2.12E+05<br>(350.5 %) |
|  | 21. Inund<sub>s</sub><br>(PFU/mL) | 6.8E+08<br>(14.6 %) | 2.95E+08<br>(74.7 %) | 6.21E+08<br>(64.5 %) | 1.11E+08<br>(36.5 %) |
|  | 22. Prolif<sub>R</sub><br>(CFU/mL) | 2.67E+06<br>(69.1 %) | 6.19E+06<br>(121.7 %) | 1.50E+07<br>(173.7 %) | 9.64E+06<br>(598.4 %) |
|  | 23. Inund<sub>R</sub><br>(PFU/mL) | 3.68E+10<br>(3.3 %) | 2.40E+10<br>(18.1 %) | 1.19E+11<br>(24.7 %) | 3.76E+09<br>(33.1 %) |

<sup>a</sup> Fixed effect parameters are as described in the main text and expressed as population estimate (relative standard error).
<sup>b</sup> Omega parameters are expressed as standard deviations of random effects (relative standard errors)
<sup>c</sup> Error model parameters are as follows: a(B), coefficient of additive error of bacteria count, a(V) and b(V), coefficient of additive and proportional error of phage count, respectively.
<sup>d</sup> Proliferation and inundation threshold values are given as mean (CV %) of Bayesian posterior estimates from each experiment.
1 to 23: Line number

**Supplementary Table S4.** List of mutated genes in SH1000 after 24 h of exposure to phage V1SA019. In bold: genes involved in the teichoic acid biosynthesis pathway.

| MOI | Mutant | Gene | Effect |
| --- | --- | --- | --- |
| 0.01 | KR19 | **tarK** | missense_variant c.536A>G p.His179Arg |
|  |  | **tarA** | missense_variant c.665G>T p.Arg222Ile |
|  |  | purR | missense_variant c.286C>T p.Arg96Cys |
|  | KR20 | S35 | intragenic_variant n.508155A>G |
|  |  | **tarK** | missense_variant c.536A>G p.His179Arg |
|  |  | tsr24 | non_coding_transcript_variant |
|  |  | **tarA** | missense_variant c.665G>T p.Arg222Ile |
|  |  | znuB | missense_variant c.143G>A p.Ser48Asn |
|  | KR21 | tsr24 | non_coding_transcript_variant |
|  |  | **tarA** | missense_variant c.665G>T p.Arg222Ile |
| 1 | KR25 | tsr24 | non_coding_transcript_variant |
|  |  | rpoB | missense_variant c.2161A>G p.Thr721Ala |
|  |  | **rfe/tagO** | frameshift_variant&missense_variant c.556delGinsAT p.Val186fs |
|  | KR26 | S35 | intragenic_variant n.508155A>G |
|  |  | tsr24 | non_coding_transcript_variant |
|  |  | mrcB | disruptive_inframe_insertion c.785_787dupGCA p.Arg262_Ile263insSer |
|  |  | **tarA** | missense_variant c.665G>T p.Arg222Ile |
|  | KR27 | mrcB | missense_variant c.691C>T p.Pro231Ser |
|  |  | **tarA** | missense_variant c.665G>T p.Arg222Ile |
| 100 | KR31 | **rfe/tagO** | deletion of 823 nucleotides |
|  | KR32 | **tarK** | missense_variant c.536A>G p.His179Arg |
|  |  | **rfe/tagO** | frameshift_variant c.853delG p.Val285fs |
|  | KR33 | **tarK** | missense_variant c.536A>G p.His179Arg |
|  |  | **rfe/tagO** | frameshift_variant c.853delG p.Val285fs |

**Supplementary Table S5.** Apparition of SH1000 resistant mutants to V1SA019 over time of kinetic according to MOI. nd: not determined.

| Resistants to V1SA019 (total tested) |  | Time of kinetic |  |  |  |  |
| --- | --- | --- | --- | --- | --- | --- |
|  |  | 8 h | 10 h | 12 h | 14 h | 24 h |
| MOI | 0.01 | 0 (30) | 0 (30) | 0 (30) | 1 (3) | 23 (24) |
|  | 1 | 1 (15) | 0 (15) | 0 (22) | 10 (10) | 30 (30) |
|  | 100 | 2 (30) | 5 (27) | nd | nd | 30 (30) |

